# Targeting OLIG2 increases therapeutic responses in SHH medulloblastoma mouse models and patient-derived medulloblastoma organoids

**DOI:** 10.1101/2022.02.14.480293

**Authors:** Yuchen Li, Taylor Dismuke, Chaemin Lim, Zara C. Bruce, Carolin Offenhäuser, Ulrich Baumgartner, Mellissa Maybury, Rochelle C. J. D’Souza, Timothy Hassall, Brandon Wainwright, Gregory Stein, Michael Piper, Terrance G. Johns, Marina Sokolsky-Papkov, Timothy R. Gershon, Bryan W. Day

## Abstract

Recurrence after therapy is the primary life-threatening complication of medulloblastoma. In Sonic Hedgehog (SHH)-subgroup medulloblastoma, OLIG2-expressing tumour stem cells are crucial to recurrence. We investigated the potential of the small-molecule OLIG2 inhibitor CT-179 to decrease recurrence in patient-derived organoids, mice genetically-engineered to develop SHH-driven MB, and mice with MB patient-derived xenograft (PDX) tumours. We found that *OLIG2* mRNA significantly correlated with poor survival in patients with SHH-MB, but not other subgroups. CT-179 rapidly downregulated OLIG2 protein *in vitro* and displayed nanomolar IC_50_ values. CT-179 arrested MB cells at G_2_/M, with degradation of cyclin B1 and phospho-CDK1 inducing apoptosis. *In vivo* CT-179 induced similar cell cycle changes in MBs in *Smo*-mutant mice and significantly increased mouse survival. In both MB organoids and mouse models, CT-179 combined with radiotherapy showed greater efficacy than either treatment alone. These data highlight the potential for OLIG2-targeted therapy to improve MB outcomes by targeting recurrent disease.

## INTRODUCTION

Brain cancer remains the leading cause of cancer-related death in children (Ostrom et al., 2018). Medulloblastoma (MB) is the most common paediatric brain cancer and accounts for approximately 20% of all cases (Ostrom et al., 2016). International consensus recognises four distinct MB molecular subgroups: Wingless (WNT), Sonic hedgehog (SHH), Group 3 and Group 4 tumours (Northcott et al., 2011). Overall survival (OS) rates have reached 70-80%, but the outcome for young children, especially infants, is worse (Gajjar et al., 2006; Rutkowski et al., 2010). Those who do survive suffer from long-term therapy-induced side effects and development of secondary tumours (Mabbott et al., 2008; Mabbott et al., 2005; Neglia et al., 2006; Packer et al., 2003; Spiegler et al., 2004). Novel therapies with specific anti-tumour efficacy and less off-target toxicity hold promise to improve both survival rates and quality of life for MB patients.

OLIG2 is a basic helix-loop-helix (bHLH) transcription factor that functions in the developing brain alternately to promote oligodendrocyte differentiation or maintain the undifferentiated state of neural progenitor and stem cells, depending on the cellular context (Meijer et al., 2012; Takebayashi et al., 2002; Zhou and Anderson, 2002). OLIG2 is also expressed and functional in a large percentage of adult glioblastoma (GBM) (Kosty et al., 2017). Recent single-cell RNA sequencing efforts have revealed a developmental hierarchy of progenitor cells in SHH-driven MB and showed that OLIG2-expressing stem-like progenitor cells are required for both MB progression and recurrence after therapy (Zhang et al., 2019) and related OLIG2+ stem cells to radioresistance (Malawsky et al., 2021). These studies suggest anti-OLIG2 therapy could be a viable tumour stem cell targeting strategy, particularly in SHH-driven disease.

A series of small molecule OLIG2 inhibitors were previously identified using a pharmacophore-guided 3D-structural search to find compounds that engage the OLIG2 dimerization interface (Tsigelny et al., 2017). This effort identified SKOG102, a compound that selectively modulated OLIG2 targets, entered the brain following IP injection, and suppressed PDX glioblastoma tumour growth *in vivo.* SKOG102 defined the initial pharmacophore for a medicinal chemistry effort that, ultimately, led to the development of CT-179. CT-179 is a novel small molecule OLIG2 inhibitor made by Curtana Pharmaceuticals (molecular weight (MW) 397.3, patent application WO2016138479A1), with favourable blood brain barrier (BBB) penetration properties. CT-179 binds to an OLIG2-DNA complex and alters OLIG2 conformation, thereby inhibiting transcriptional activity. CT-179 was recently awarded FDA Rare Paediatric Disease Designation for the treatment of MB and is administered via a tablet formulation.

The present effort sought to assess the levels of OLIG2 expression in MB subgroups and to evaluate the efficacy of CT-179 as a single agent and in combination with radiotherapy (RT) in MB cell lines, MB explant tissue organoids (MBOs), genetically-engineered mouse (GEM) model and patient-derived xenograft (PDX) models. Analysis revealed a significant correlation between *OLIG2* mRNA expression and shorter survival in SHH subgroup patients, but not other subgroups. We detected elevated *OLIG2* mRNA and protein expression in several primary MB cell lines and PDX models irrespective of MB subgroup. CT-179 showed significant *in vitro* and *in vivo* anti-tumour efficacy, both as a single agent and when combined with RT in SHH-driven disease. Our findings validate the potential of targeting OLIG2 in SHH-MB and highlight the novel small molecule CT-179 for further clinical evaluation, particularly when combined with RT in patients with MB.

## RESULTS

### OLIG2 correlates with poor survival in SHH-MB patients

To investigate whether *OLIG2* mRNA expression correlated with patient survival, we analysed the Cavalli MB dataset (Cavalli et al., 2017). Results show elevated *OLIG2* significantly correlated with poor survival in SHH-MB patients, but not other subgroups (Figure 1A). These findings indicate a potential functional link between OLIG2 and SHH-MB tumourigenesis. Next, we confirmed *OLIG2* mRNA expression in a panel of immortalized MB lines, primary MB lines and PDX models obtained from the Olsen laboratory and generated in-house (R201, R203 and R302) (Brabetz et al., 2018). Results show *OLIG2* was generally higher in cell lines in comparison to PDX tumours (Figure 1B). Elevated OLIG2 protein was confirmed in our panel of MB cell lines (Figure 1C). In PDX tumours from the Olsen group, OLIG2 expression was higher in the SHH Med-813 model compared to the Group 3 Neo-113 model, which was consistent with the gene expression results analysed from the R2 database (Figure 1D). In addition, immunohistochemistry (IHC) staining for OLIG2, performed on MB PDX tumours, showed positive OLIG2 focal staining (Figure 1E).

**Figure 1.**
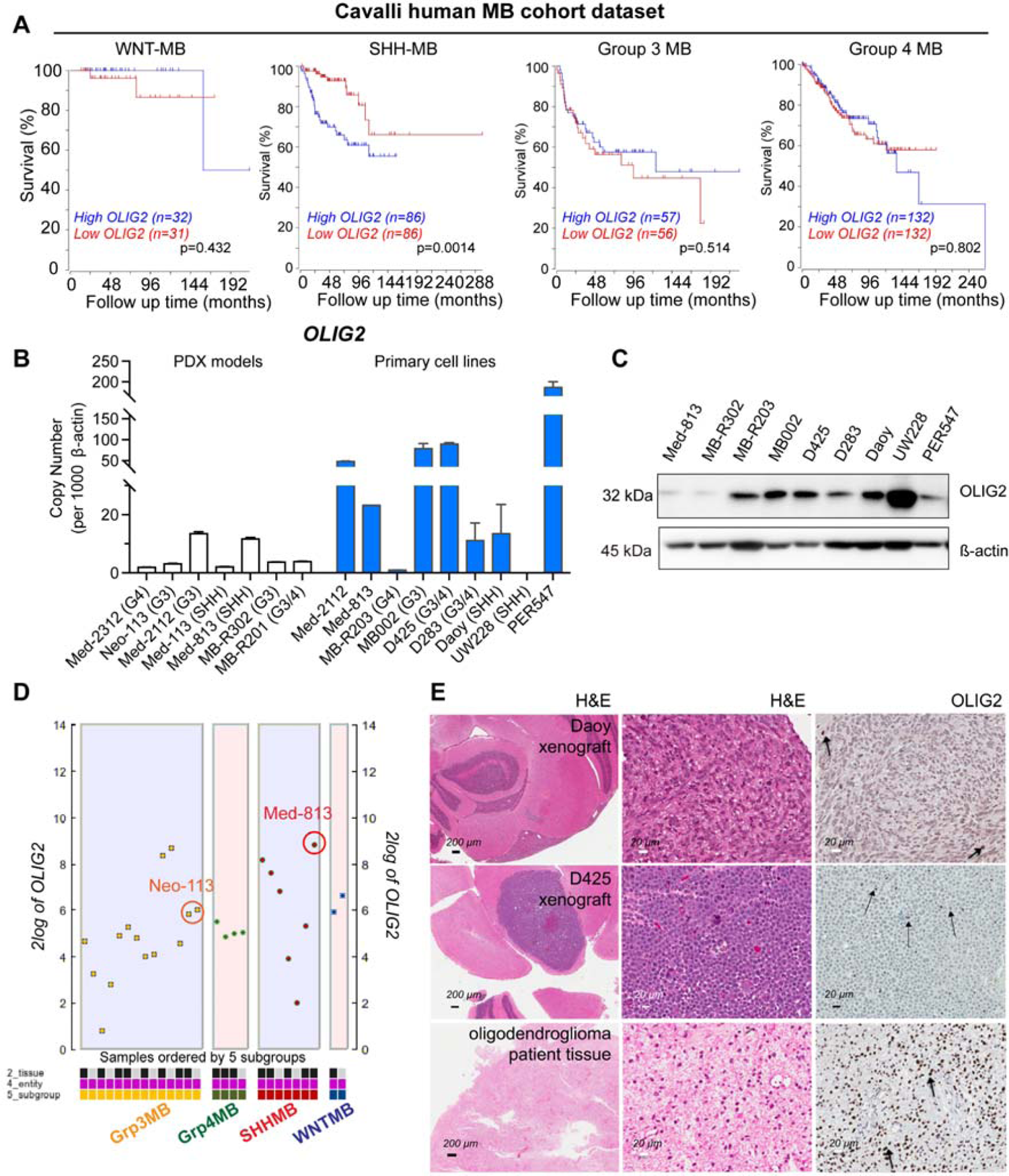
OLIG2 expression in paediatric MB. (A) High *OLIG2* expression correlates with poor OS in SHH-MB, with a subgroup specific manner. Kaplan-Meier curves of four subgroups of MB patients based on OLIG2 expression. (B) *OLIG2* expression in PDX models, primary cell lines and ATCC paediatric MB cell lines (data are shown as means ± SD, n=6). (C) OLIG2 expression at protein level in MB cell lines. (D) *OLIG2* expression of Med-813 and Neo-113 filtered tissues were analysed using R2: Genomics analysis and visualization platform. https://hgserver1.amc.nl/cgi-bin/r2/main.cgi. (E) OLIG2 immunostaining in 2 MB xenograft models Daoy and D425. Oligodendroglioma patient tissue was used as a positive control. Arrowheads, OLIG2^+^ cells. Scale bars are indicated in (E).

### CT-179 induces apoptosis

CT-179 is an orally bioavailable OLIG2 small molecule inhibitor (MW: 397) (Figure 2A). Initially, Daoy, UW228 and Med-813 MB cells were treated with a serial dilution of CT-179 for 7 days. All lines tested displayed nanomolar activity (Figure 2B). A reduction in OLIG2 protein was observed 48 hours post CT-179 treatment, demonstrating effective on-target specificity (Figure 2C and D). Apoptosis analysis showed CT-179 (1 µM) alone induced significant cell death at three days, which was further enhanced when used in combination with RT (2Gy) (Figure 2E and F). We noted this response matched the kinetics of reduced OLIG2 following CT-179 therapy. Reduced expression of cleaved poly-ADP ribose polymerase (PARP) coincided with increasing cleaved caspase 3 overtime, indicating a robust apoptotic response (Figure 2G). Furthermore, CT-179 led to a reduction in expression of the anti-apoptotic proteins Mcl-1, Bcl-2, and Bcl-xL (Figure 2H).

**Figure 2.**
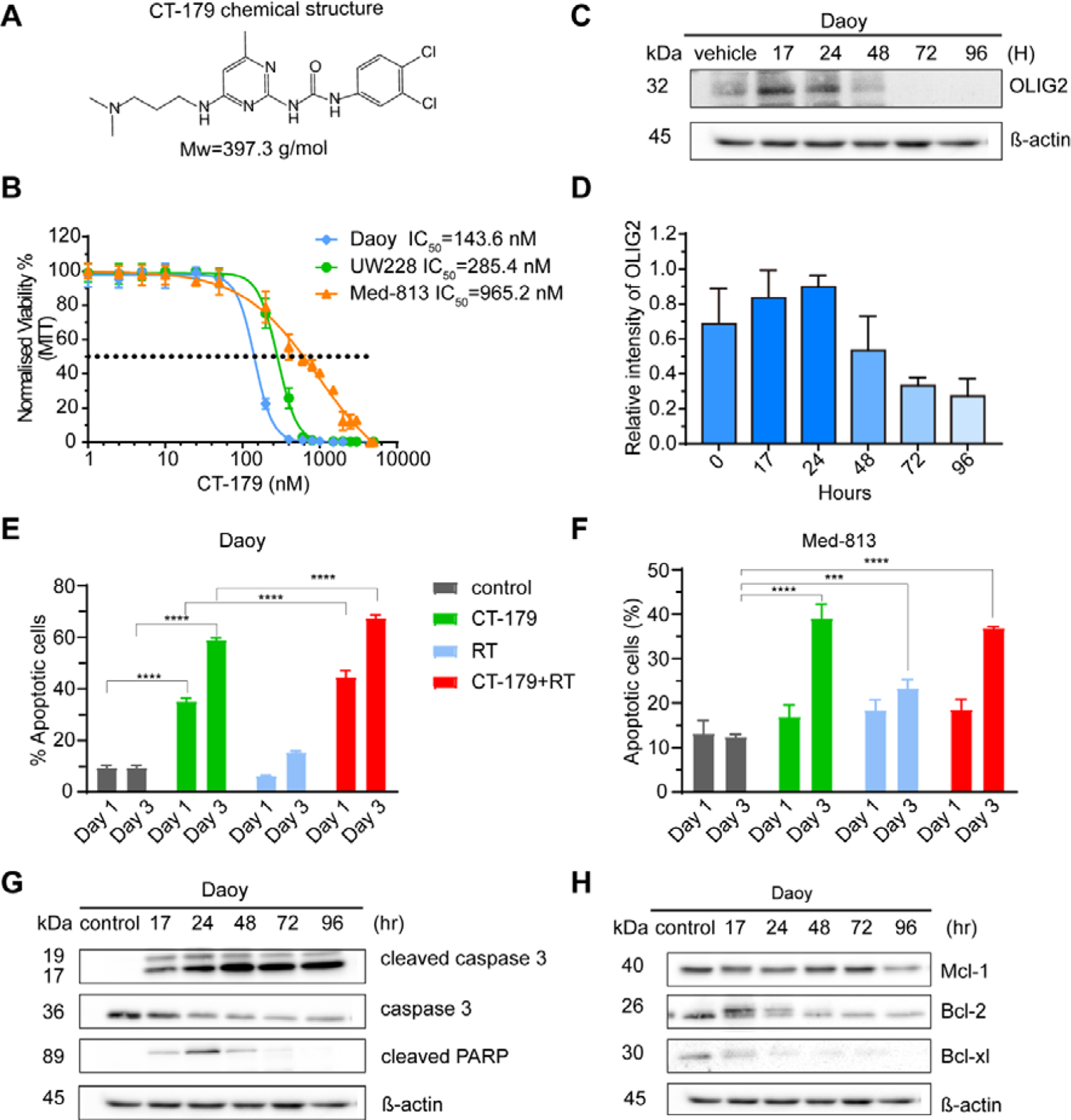
CT-179 induces MB cell apoptosis. (A) Chemical structure of CT-179. (B) MB cell lines were treated with CT-179 for 7 days and a median inhibitory concentration (IC_50_) determined experimentally for each cell line (data are shown as means ± SD, n=3). (C) Daoy were treated with 1 µM CT-179 for 17, 24, 72 or 96 hours, after which total protein were analysed. OLIG2 expression peaked at 17 and 24 hours after treatment, then decreased to basal and completely diminished after 72 hours. (D) Quantitation of OLIG2 expression in Daoy in response of CT-179 (data are shown as means ± SD, n=2). (E and F) Percentage of Annexin V^+^ cells in Daoy cells and Med-813 treated with CT-179 (1 µM) or RT (2 Gy) alone, or in combination (means ± SD, ***p < 0.001 ****p < 0.0001, n=6). (G) Representative western blots show expression of cleaved caspase-3, caspase 3 and cleaved PARP in Daoy after treated with CT-179 (1 µM). (H) Representative western blots show expression of Mcl-1, Bcl-1 and Bcl-xL in Daoy after treated with CT-179 (1 µM).

### CT-179 induces mitotic arrest

Lu and colleagues had previously shown that OLIG2 could regulate multiple cell cycle-associated transcriptional regulators (Lu et al., 2016). Further studies highlighted the ability of OLIG2 to regulate genes controlling the cell cycle and microtubule function during developmental (Darr et al., 2017). We therefore, assessed the cell cycle following CT-179 (1µM) treatment alone and in combination with RT (2Gy). Results show a G_2_/M phase arrest following single therapy, which was further pronounced when combined with RT (Figure 3A). Molecular analysis showed that CT-179 disrupted the orderly progression of key mitotic mechanisms, demonstrated by the altered expression of cyclin B, Cyclin Dependent Kinase 1 (CDK1), phosphorylated CDK1 (p-CDK1) and phosphorylated Histone H3 (p-HH3). Mitotic cells typically express cyclin B1 at the onset of mitosis when bound CDK1 mediates spindle assembly and mitotic entry (Sullivan and Morgan, 2007), then rapidly degrade cyclin B1 after the spindle assembly checkpoint. Simultaneously, cells typically inactivate CDK1 by dephosphorylation and pHH3 which resolves following telophase (Sullivan and Morgan, 2007). CT-179 treatment decreased cyclin B1, CDK1 and p-CDK1, while maintaining persistently elevated pHH3 (Figure 3B). Med-813 responded to CT-179 with a more complex disruption of mitotic mechanisms, with initial increase in cyclin B1, CDK1 and p-HH3 over 24 hours with decreased p-CDK1, followed by a decrease in cyclin B1 and CDK1 and a marked increase in p-CDK1 at later time points (Figure 3D). To explore the effects of CT-179 on nuclear morphology and mitotic spindle formation, Hoechst (cyan) staining of the α-tubulin (yellow) staining of mitotic spindles was conducted. Immunofluorescence (IF) staining results illustrated cells treated with vehicle had a normal nuclear morphology and spindle alignment, whereas cells treated with CT-179 took on a multinucleated/tetraploid appearance, with satellite micronuclei (white arrow) and an ancillary nuclear lobe formation (green arrow) (Figure 3C). These studies show that CT-179 treatment resulted in profound mitotic disruption.

**Figure 3.**
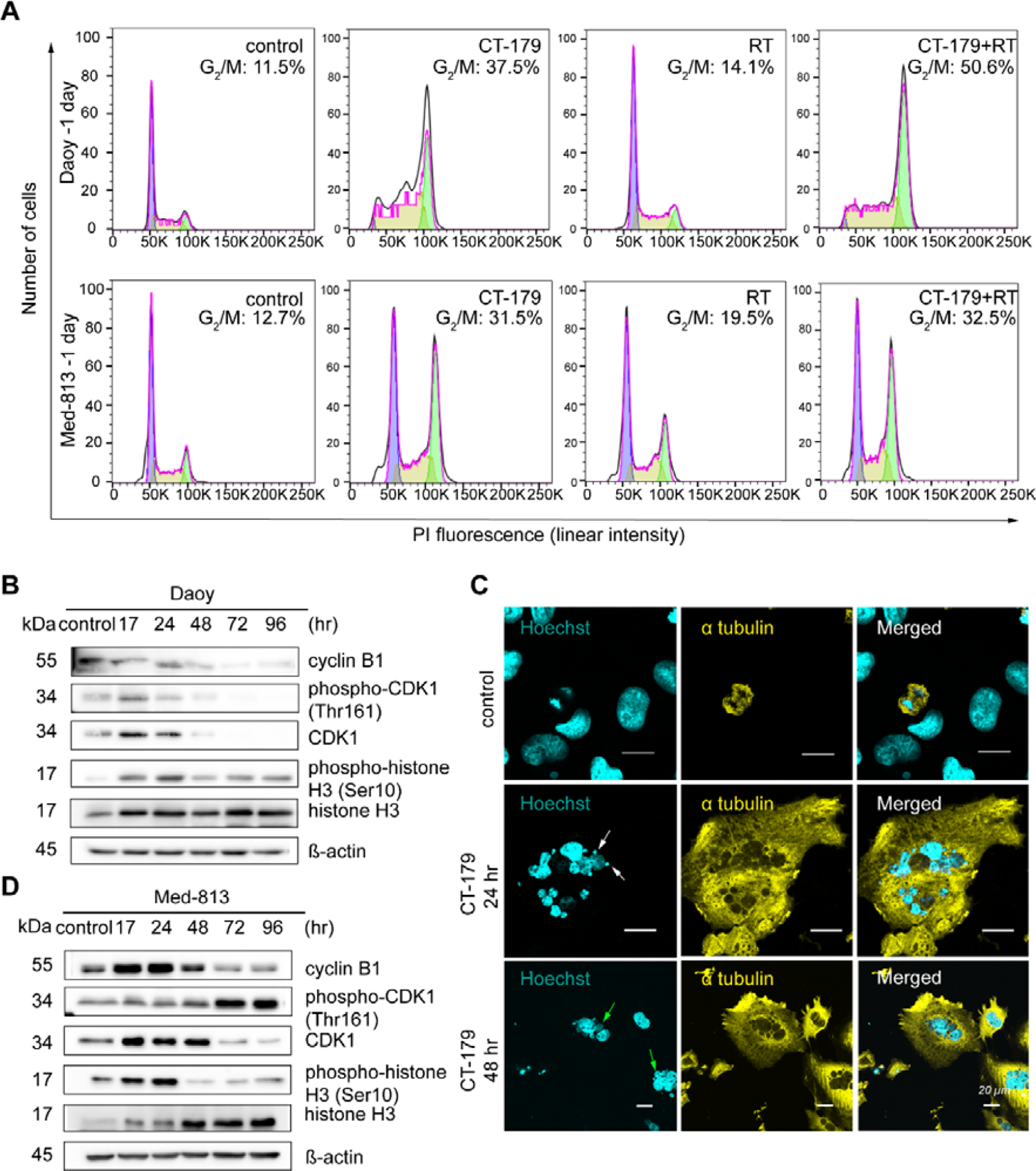
CT-179 induces apoptosis and mitotic slippage in MB cells. (A) Histograms of Daoy and Med-813 cells fixed and stained with propidium iodide after 24 hours treatment with vehicle (left panels) or 1 µM CT-179 or RT (2 Gy) alone, or in combination. CT-179 has synergistic anti-tumour effect when used in combination with RT. (B) Representative western blots show expression of cyclin B1, phospho-CDK1 and total CDK1 in Daoy after treated with CT-179 (1 µM). (C) Top panel shows a Daoy cell in anaphase with normal nucleus morphology and spindle alignment. CT-179 treatment (1 µM) results in abnormal nucleus phenotypes including satellite micronuclei (white arrow) and ancillary nucleus lobe formation (green arrow), suggesting cells undergo a mitotic slippage response. (D) Representative western blots show expression of cyclin B1, phospho-CDK1 and total CDK1 in Med-813 cells after CT-179 (1 µM) treatment.

### CT-179 treatment induces cell death in MB explant organoids (MBOs)

Tumour explant organoids are generated directly from tumours surgically resected from patients without cell dissociation. This approach allows tumour stroma, blood vessels and immune infiltrate to remain intact. We adopted a recent adult GBM explant approach (Jacob et al., 2020a; Jacob et al., 2020b) to freshly resected MB specimens. To the best of our knowledge, we are the first group to adopt this technique to medulloblastoma, generating MBOs that could be cultured for up to 12 weeks. DNA methylome profiling classified three selected specimens, R403 (Group 4-MB, subtype VI), R901 (Group 4-MB, subtype VIII) and R902 (SHH-MB). With limitation of the tissue size, we were only able to assess *OLIG2* in two of the tumours, R901 and R902; both had detectable *OLIG2* expression that was greater in R902 (Figure 4A).

**Figure 4.**
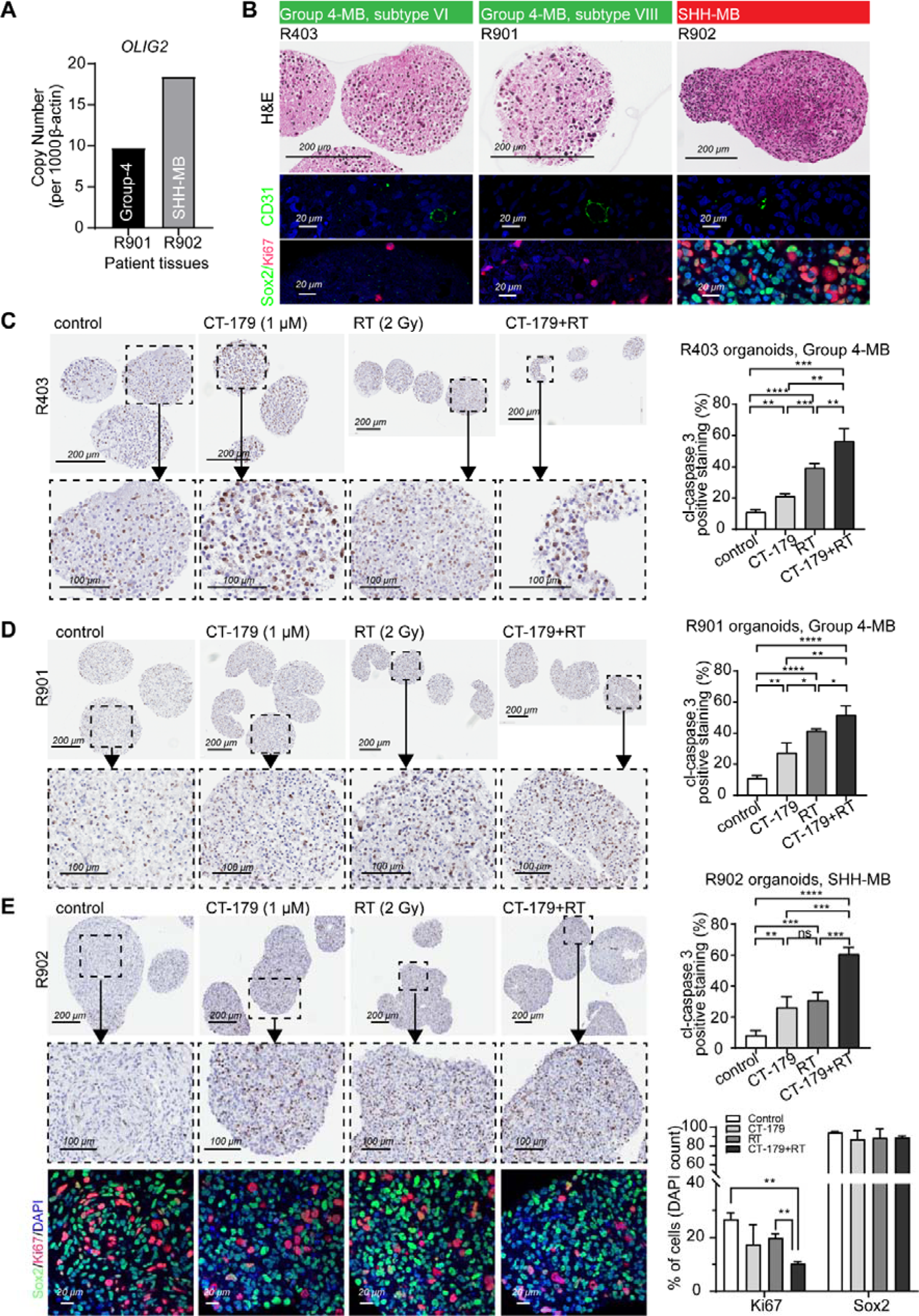
CT-179 induces apoptosis in MB explant organoids R403, R901 and R902. (A) *OLIG2* expression in patient tissues R901 and R902 (means ± SD, n=2). (B) H&E of untreated MB organoids R403, R901 and R902. CD31 stained for endothelial cells. Ki67 stained for proliferating cells. Sox2 is highly expressed in SHH MB, in both glial cells and tumour stem cells. (C) R403, (D) R901 and (E) R902 were treated for CT-179 (1 μM) or RT (2 Gy) alone or in combination for 48 hours. Cells were stained with cleaved caspase 3 (cl-caspase 3), Ki67 and Sox2 (means ± SD, n=3, *p < 0.05, **p < 0.01, ***p < 0.001, ****p < 0.0001). Combination treatment significantly induced cell death in SHH-MB organoids compared to single treatment.

The H&E and IF results show that viable MBOs formed with heterogeneous morphology, intact blood vessels (CD31+) and molecular features (Sox2+ in SHH-MB) (Figure 4B) (Ahlfeld et al., 2013). R403, R901 and R902 MBOs were treated with CT-179 (1 µM), RT (2 Gy), or combined for 48 hours. Cleaved caspase 3 staining showed the R403 and R902 MBOs were the most sensitive to CT-179 alone and displayed significant cell death following therapy (Figure 4C). Combination treatment affected the structure of the R403 organoids resulting in gross MBO collapse. The R901 organoid responded similarly to therapy while the MBO structure was more stable (Figure 4D). The response of the SHH subgroup R902 organoid to combination treatment was more pronounced than the group 4 organoids (Figure 4E). Comparing the effects of treatment on the fractions of cells in the R902 MBOs that expressed the proliferation marker Ki67 and the glial/stem cell marker SOX2 (Ocasio et al., 2019; Vanner et al., 2014) demonstrated that combination therapy significantly reduced proliferation without inducing a compensatory increase in the SOX2+ population (Figure 4E).

### In vivo pharmacokinetic studies of CT-179 in mice

Two independent CT-179 pharmacokinetic (PK) animal studies, including brain uptake, were performed by Biodura Inc. The initial study (EXT-240) assessed the bioavailability and concentration versus time of CT-179 in mice administered either a single intravenous (IV) (1mg/Kg) or oral dose (per os, PO) (20mg/Kg) (Figure S1A and B). Three mice (C57Bl/J6) were tested in each of the IV and PO groups. Results highlighted consistent plasma concentrations, which were maintained across a 24-hour period, indicating a suitable half-life for once per day dosing (Figure S1C). Increasing CNS exposure were detected up to 4 hours post dose, with an estimated brain to plasma (B/P) ratio reaching an average of 6.72. While the B/P ratio was well above one, an equilibrium was achieved and the brain tissue concentration did not continue to rise with prolonged exposure. When CT-179 was discontinued, the drug concentration in the brain decreased over time (Tsigelny et al., 2017). CT-179 displayed a predicted half-life in the range of 10-12 hours and high bioavailability (range estimated as 58-91%; mean 75%) (Figure S1D). The objective of the second study (EXT-241) was to estimate steady state exposures of CT-179 in mice receiving PO doses of 1 and 5 mg/Kg (Figure S1E). Results show both PO doses achieved significant plasma levels. CT-179 exposures were measured at 24 hours on Days 1 and 3 to compare the initial dose to steady state levels. Findings indicate that CT-179 levels increased approximately 35% between the two time points. CT-179 displayed an estimated brain to plasma ratio of >10 for both 1 and 5 mg/Kg groups, demonstrating CNS penetration and accumulation (Figure S1F and G). Plasma and CNS exposure diminished 48 hours after the last dose implying that equilibration of the compound is reversible (Figure S1G). Taken together, the PK studies demonstrated that CT-179 had high oral bioavailability and achieved measureable plasma and brain exposures over 24 hours with somewhat increased exposure between Days 1 and 3 following once daily dosing.

### CT-179 disrupts OLIG2 processing and cell cycle progression in SHH-driven tumours

We generated MB-prone mice by crossing *hGFAP-Cre* and *SmoM2* mouse lines to produce *hGFAP-Cre/SmoM2* (*G-Smo*) pups. These animals developed SHH-MB tumours with 100% frequency by P10. CT-179 was administered via intraperitoneal (IP) injection (50-150 mg/kg) with a maximum tolerated dose (MTD) of 80 mg/kg every other day (EOD) or 100 mg/kg every three days. Higher doses were associated with instances of brain haemorrhage (Figure S2), however no haemorrhage was observed at 80 mg/kg EOD or 100 mg/kg every three days. We next administered CT-179 80 mg/kg at P10, P12, P14 and P16 and assessed phosphorylated OLIG2 (pOLIG2) levels 24 hours post therapy. Results show a decrease in pOLIG2, accompanied by reduced phosphorylated RB (pRB) (Figure 5A). These findings confirmed the on-target specificity of CT-179 *in vivo*.

**Figure 5.**
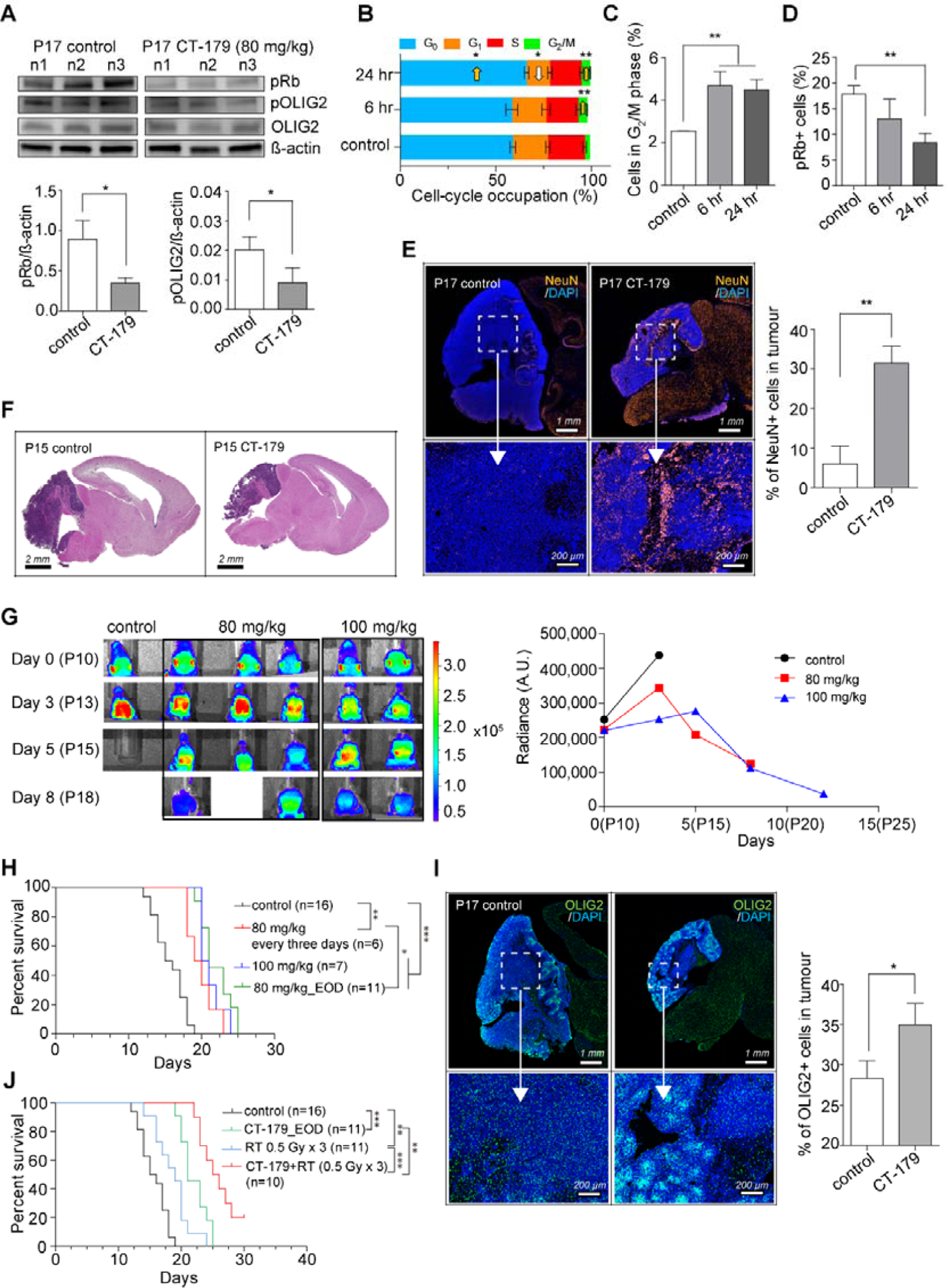
CT-179 efficacy in *G-Smo* model. (A) Western blots for expression of pRb and pOLIG2 in G-*Smo* MB 24 hours after CT-179 treatment (EOD, 80 mg/kg from P10, harvested at P17). Quantification of pRb and pOLIG2. The level of expression was normalized by β-actin (means ± SD, n=3, *p < 0.05). (B) Cell-cycle occupation analysis of 80 mg/kg of CT-179 treated (6 and 24 hours) *G-Smo* mice (means ± SD, n=3). (C and D) Quantification of (C) G_2_/M phase and (D) G_1_ cells showing highly phosphorylated Rb in CT-179-treated (6 and 24 hours) mice (means ± SD, n=3, **p < 0.01). (E) Representative NeuN/DAPI IF in *G-Smo* MB treated with CT-179 100 mg/kg or saline every three days, 24 hours after the final treatment with quantification of NeuN+ cells in tumours of replicate mice (means ± SD, n=3, **p < 0.01). (F) H&E of CT-179 treated brains at P15 (EOD, 80mg/kg from P10) compared to saline controls. (G) Longitudinal dynamic tracing of CT-179 in luciferase-tumour bearing mice treated as indicated every three days. Quantified in the right panel. (H) Kaplan-Meier curves of *G-Smo* mice on the indicated monotherapy regimens (*p < 0.05, **p < 0.01, ***p < 0.001). (I) Representative OLIG2/DAPI IF in *G-Smo* MB treated with CT-179 (80 mg/kg) or vehicle EOD and harvested at P17, 24 hours after final dose, with quantification of OLIG2 IF in replicate mice (means ± SD, n=3, *p < 0.05). (J) Kaplan-Meier curves of *G-Smo* mice on the indicated treatment regimens (**p < 0.01, ***p < 0.001

To assess cell cycle effects, tumours were harvested 2, 6 and 24 hours post CT-179 (80 mg/kg) therapy. Animals were injected with EdU 40 mg/kg, IP 30 minutes prior to harvest. We dissociated tumours, stained for EdU, phosphorylated RB (pRB) and DNA content, and quantified each marker by flow cytometry, gating as in Figure S3. At 6 hours, CT-179-treated tumours showed significantly increased G_2_/M fractions, consistent with our *in vitro* data. By 24 hours, G_2_/M remained increased, G_0_ also increased and G_1_ decreased, indicating G_0_ and G_2_ arrest (Figure 5B and C). Consistent with reduced cycling, the pRB+ population decreased by 24 hours (Figure 5D).

Analysis of tumour pathology showed increased neuronal differentiation in CT-179 treated tumours, demonstrated by increased NeuN expression (Figure 5E), consistent with cell-cycle exit. Increased differentiation correlated with reduced growth, as CT-179-treated animals consistently displayed smaller tumours at P15 (Figure 5F). These *in vivo* pharmacodynamic effects show that CT-179 effectively penetrated the blood brain barrier at bioactive concentrations and increased differentiation along regionally appropriate developmental trajectory.

### CT-179 reduces tumour growth in G-Smo mice with SHH-MB

To assess the *in vivo* efficacy of CT-179, we first crossed *Gli-luc* mice that carry a GLI-activated luciferase reporter transgene (Becher et al., 2008) into the *G-Smo* breeders to generate *G-Smo^Gli-luc^* pups. We then compared bioluminescence imaging (BLI) in *G-Smo^Gli-luc^* mice treated with CT-179 or vehicle, administered every three days. Control *G-Smo^Gli-luc^* mice showed an increasing BLI signal over time, reflecting cumulative tumour growth with SHH hyperactivation (Figure 5G control). In comparison, mice treated with CT-179 showed a dose-dependent decrease in BLI signal (Figure 5G).

To determine if CT-179 produced clinically relevant tumour suppression, we analysed the event-free survival time (EFS) of *G-Smo* mice until symptomatic tumour progression, comparing CT-179 treatment to vehicle-treated controls. Three regimens were tested, CT-179, IP: 80 mg/kg EOD, 80 mg/kg every three days, or 100 mg/kg every three days. Treatment commenced at P10, when tumours were detectable. Treatment continued until progression. All three CT-179 regimens significantly improved EFS compared to control (Figure 5H). Analysis of tumours at P17 showed that the fraction of OLIG2+ cells increased in mice treated with CT-179 from P10-P17, suggesting a homeostatic response to OLIG2-targeted therapy may mediate resistance in single-agent treatment (Figure 5I).

To assess efficacy of CT-179 when combined with RT, we treated *G-Smo* mice with RT combined CT-179 and compared to RT alone, CT-179 alone, and vehicle-treated controls (Figure 5J). We noted potential for combinatorial toxicity, as initial studies combining 80 mg/kg EOD CT-179 with 5 fractions of 2 Gy RT resulted in mouse deaths earlier than expected from tumour progression, and we accordingly adjusted the radiation dose downward (Table S1). We found that 3 fractions of 0.5 Gy was tolerable and produced a statistically significant therapeutic effect, with *G-Smo* mice treated with 3 fractions of 0.5 Gy showing longer EFS compared to untreated controls (Figure 5J and Table S2). The combination of CT-179 plus RT significantly improved EFS compared to either treatment alone (Figure 5J). The addition of CT-179 thus increased the efficacy of RT in a mouse model of highly aggressive and treatment-refractory SHH MB.

### CT-179 synergises with radiotherapy in orthotopic xenograft models

To determine if CT-179 showed similar clinically relevant efficacy against human MB, and to determine efficacy when combined with RT, we orthotopically (cerebellum) engrafted Daoy-luci (SHH-MB) and PDX models Med-813-luci (SHH-MB) and Neo-113 (Group 3-MB) into immune-compromised NOD *rag* gamma (NRG) mice. Tumour formation was confirmed by BLI. Four treatment arms were assessed: vehicle control, CT-179 (50mg/kg, IP twice weekly for two weeks), RT (8Gy in 2Gy fractions) and treatment combined (Figure 6A). In the Daoy model, one mouse in the CT-179 single treatment arm had skin irritation at the injection site and was censored on Day 28 (Figure 6B and C). Another showed swelling of the abdomen at the injection site and was censored on Day 35 (Figure 6B and C). Animals receiving CT-179 + RT treatment progressed noticeably slower, with significantly delayed tumour growth. No tumours were detected via BLI in three animals in the combination group 36 days post commencement of treatment (Figure 6B). CT-179 combined with RT led to a significant increase in EFS when compared to vehicle control, median survival: vehicle control (55 days), CT-179 (60 days) or RT (61.5 days) or combination (75.5 days) (Figure 6C).

**Figure 6.**
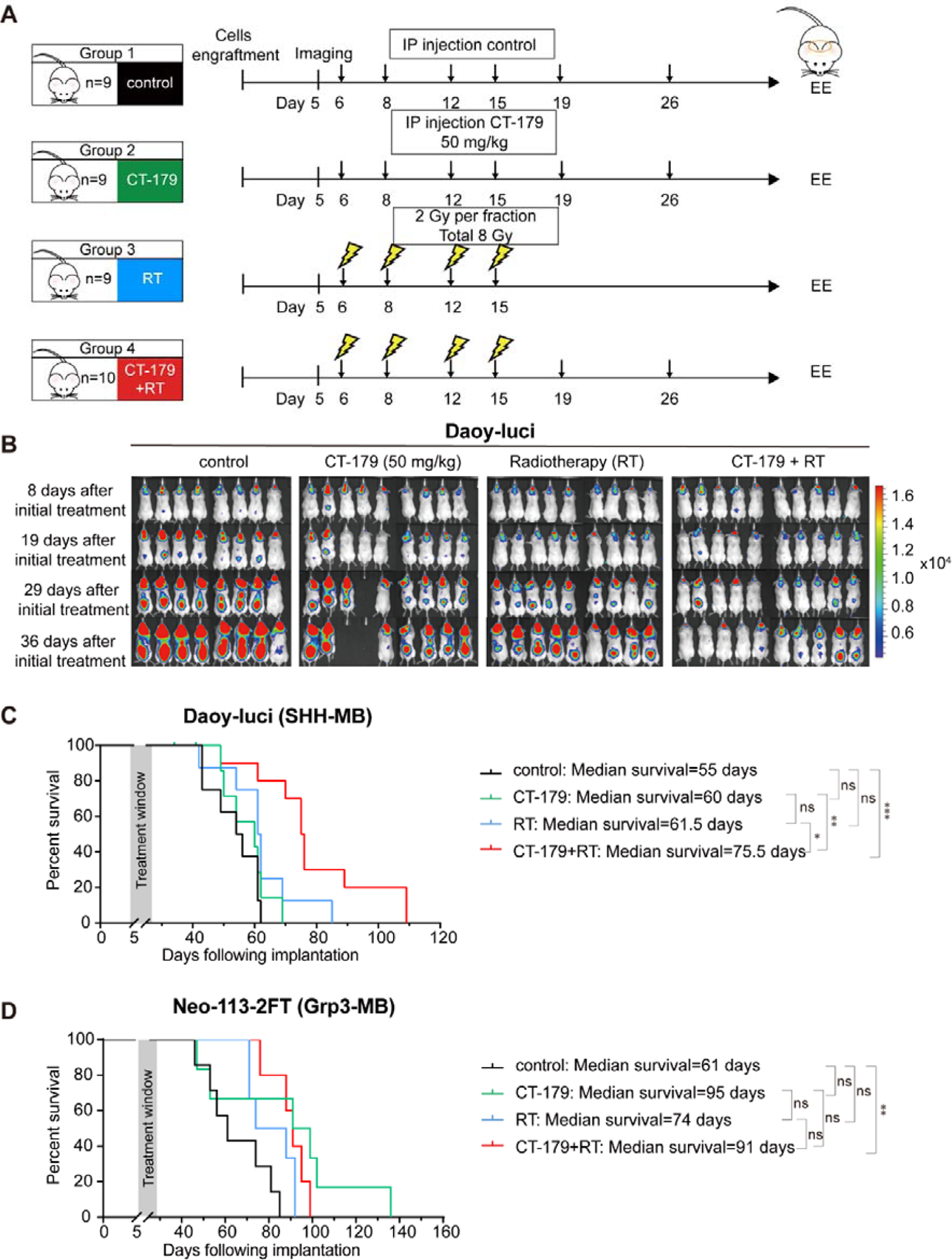
CT-179 administered IP exerts synergistic anti-tumour activity when used in combination with RT in orthotopic MB models. (A) Treatment plans used in xenograft animal models Daoy-luci and Neo-113. (B) Representative *in vivo* bioluminescence imaging of NOD *rag* gamma mice engrafted with 1 x 10^5^ Daoy-luci cells treated with CT-179 or RT alone, or in combination. (C and D) Kaplan-Meier survival curves of Daoy-luci mice (≥ 6 per group) and Neo-113 (≥ 4 per group) treated with indicated therapies (*p < 0.05, **p < 0.01, ***p < 0.001).

Given these promising results, we progressed to assessing the primary PDX models Med-813-luci (SHH-MB) and Neo-113 (Group 3-MB) using oral gavage, to mimic an oral tablet formulation regime. Neo-113 engrafted animals were treated with the same regimen as the previous Daoy model. Log Rank test and Kaplan-Meier survival curves show CT-179 combined with RT significantly prolonged EFS compared to vehicle, while single agent treatment did not result in statistically significant changes (91 days) (**p < 0.01, Figure 6D).

We next treated Med-813 (SHH-MB) engrafted animals with 75 mg/kg CT-179 via oral gavage EOD (Figure 7A). With this regimen, CT-179 + RT significantly increased EFS (100.5 days) compared to vehicle (71 days, ***p < 0.001), CT-179 alone (76 days, *p < 0.05) and RT alone (88.5 days, *p < 0.05) (Figure 7B, C and S7). The prolonged EFS with CT-179 + RT compared to RT alone indicating the ability of CT-179 to increase radiation sensitivity. This finding was consistent with the previously reported role of OLIG2+ cells in radiation resistance (Malawsky et al., 2021; Zhang et al., 2019).

**Figure 7.**
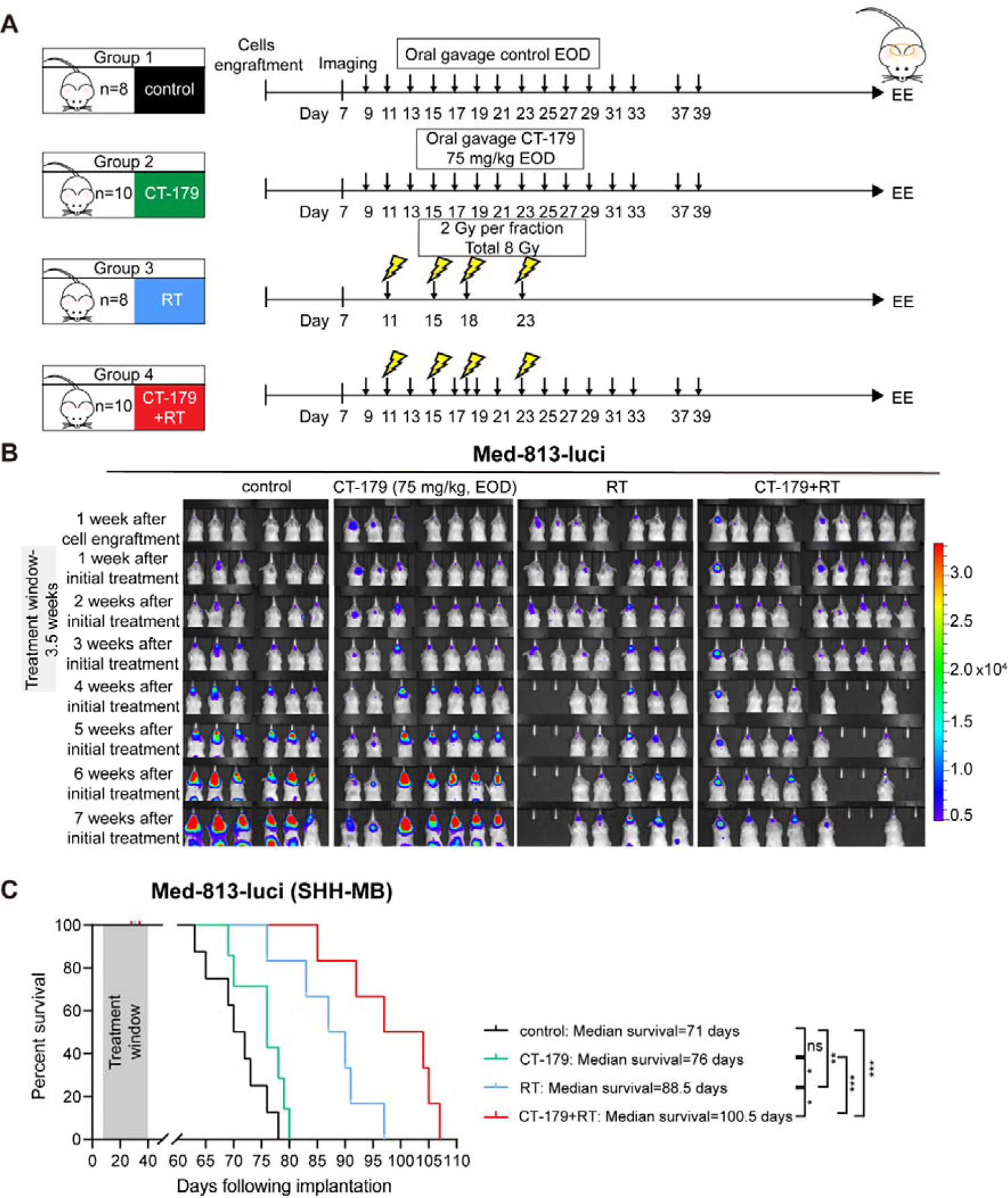
CT-179 administered via oral gavage exerts synergistic anti-tumour activity when used in combination with RT in Med813-luci model. (A) Treatment plans used in xenograft animal models Med813-luci. (B) *In vivo* bioluminescence imaging of NOD *rag* gamma mice engrafted with 3 x 10^5^ Med813-luci cells treated with CT-179 or RT alone, or in combination. (C) Kaplan-Meier survival curves of Med-813-luci mice (≥ 6 per group) treated with indicated therapies (*p < 0.05, **p < 0.01, ***p < 0.001).

We noted toxicities in both the RT group and the CT-179 + RT group. Two animals from the RT group and three animals from the combination group were censored due to side effects. These mice displayed weight loss (Figure S4A and B) and enlarged spleens (Figure S5A and B) which were also observed in the toxicity study. Animals presented with slight pallor and hunched posture. Histopathological spleen abnormalities, decreased red blood cell (RBC) count, haemoglobin (Hgb), and platelets (Plt) were noted in CT-179 + RT group (Figure S6).

## DISCUSSION

Paediatric brain cancer accounts for significant morbidity and mortality among childhood cancer sufferers. MB requires aggressive radiotherapy and chemotherapy to minimize recurrence, but at the cost of long-term cognitive, psychosocial and medical complications. Novel therapies are needed to reduce recurrence and lessen disease burden. In this study, we demonstrate that the OLIG2 inhibitor CT-179 can increase the efficacy of multi-modal therapy, most prominently in SHH-driven MB, potentially enabling new regimens that prevent recurrence while causing less neurologic and systemic toxicity.

Many lines of evidence suggest the potential efficacy of targeting OLIG2-expressing cells in MB. OLIG2 is a transcription factor that is expressed in brain progenitors during normal development and in MB stem cells (Zhang et al., 2019). OLIG2-expressing MB cells divide rapidly during the initial phase of tumourigenesis and then become a quiescent reservoir of stem cells that can drive recurrence after therapy (Zhang et al., 2019). Studies of OLIG2 protein expression in MB show foci of OLIG2 positive cells in 75% of cases (Ligon et al., 2004; Schuller et al., 2008). Both classic MB (40%) and desmoplastic MB (77%) histological subgroups exhibit OLIG2 positivity (Schuller et al., 2008). Our analysis of *OLIG2* mRNA expression and survival times in MB patients demonstrated a correlation between OLIG2 expression and recurrence risk that was specific to the SHH subgroup. These data support OLIG2 targeting as a new approach to MB therapy, particularly for SHH subgroup patients.

We studied CT-179 pharmacodynamics and efficacy *in vitro* using cell lines and clinically relevant explant MB organoids, and *in vivo* using GEMM and PDX models. *In vitro*, CT-179-treated MB cells accumulated in G_2_/M phase and underwent significant apoptosis which correlated with down regulation of OLIG2 protein levels CT-179 drug treatment. CT-179 induced gradual degradation of cyclin B1 and phospho-CDK1 and caused cells to exit mitosis without dividing chromosomes into anaphase. Consistent with our IF staining results, tetraploid MB cells were observed after treatment that exited mitosis through mitotic slippage (Galimberti et al., 2011). Protein expression analysis demonstrated accumulation of cleaved PARP 24 hours post treatment onwards, indicating rapid and significant cell death *in vitro*.

We generated, MB tumour explant organoids (MBOs) to recapitulate key-aspects of tumour biology *in vitro*. Recently Jacob and colleagues reported the generation and biobanking of patient-derived GBM 3D explant organoids, known as GBOs (Jacob et al., 2020a; Jacob et al., 2020b). Most 2D and 3D *in vitro* model systems do not preserve the tumour microenvironment, blood vessel structure and immune infiltrate due to tissue digestion and cell dissociation. GBOs are generated with minimal perturbation, fully maintaining heterogeneity for up to 12 weeks. The utility and clinical relevance of GBOs is increasingly being recognised (Jacob et al., 2020b). To the best of our knowledge, we are the first group to have successfully generated MBOs using the Jacob et al approach. We generated three MBOs which displayed morphology faithful to the original tumour. Subsequent testing with CT-179 alone or in combination with radiation showed increased percentages of cleaved caspase 3-positive cells and decreased Ki67-positive cells providing evidence of efficacy in a relevant model-based system of patient-derived MB tumour.

We used the *G-Smo* GEMM model of SHH-MB to study CT-179 efficacy *in vivo*. This model features an intact BBB (Phoenix et al., 2016) and cellular heterogeneity that resembles SHH-MB in patients (Riemondy et al., 2021). However, unlike most MBs that occur in patients, the *G-Smo* tumours are refractory to conventional therapy (Malawsky et al., 2021). In *G-Smo* mice, CT-179 showed pharmacodynamic effects that were consistent with *in vitro* studies and demonstrated effective targeting of OLIG2. CT-179 increased G_0_ and G_2_/M populations in *G-Smo* tumours, and induced terminal differentiation, demonstrated by decreased Rb phosphorylation and increased NEUN+ populations. CT-179 treatment slowed tumour progression, demonstrated by BLI in *G-Smo^Gli-luc^* mice, and prolonged survival of mice with MB.

All mice eventually progressed on single-agent CT-179 therapy, indicating the need for CT-179 to be combined with additional therapeutic modalities. The combination of CT-179 plus RT was more effective than either treatment alone, indicating that in the radioresistant *G-Smo* model, CT-179 was able to enhance RT efficacy and forestall recurrence.

Orthotopic xenograft and PDX models enabled *in vivo* testing of CT-179 efficacy in human MB. To mimic an oral regime, CT-179 was administered by gavage in our primary PDX model experiments. In the PDX models, we noted that CT-179 prolonged survival with tolerable toxicity when used in combination with RT.

Consistent with a specific role for OLIG2 in SHH-MB, CT-179 produced greater responses in the SHH-driven Daoy and Med-813 than in the Group 3 Neo-113 model. This difference in efficacy was expected based upon the negative survival correlation shown for OLIG2 in patients with SHH-MB. However, OLIG2-expressing cells are also found in other subgroups indicating that OLIG2 may contribute to tumour heterogeneity across MB disease subsets (Cavalli et al., 2017), which may explain the modest response observed in the Group 3 Neo-113 model. Together our studies in GEMM and PDX models demonstrate significant potential for CT-179 to enhance MB therapy in patients.

SHH-MB tumours constitute the pre-dominant tumour type in young children (<3 years of age) and adults (>17 years of age) (Hovestadt et al., 2020; Northcott et al., 2011). The <3 years of age cohort, in-particular, are at a stage of significant physiological and neurological development. Oligodendrocyte and myelin development (myelination) are critical for complex forms of network integration and are associated with broad aspects of higher brain function (Chen et al., 2020). OLIG2 marks proliferating, differentiation, and mature oligodendrocytes, and plays an important role in myelination (Maire et al., 2010). Myelin deficits caused by OLIG2 deficiency could lead to cognitive dysfunction and increased vulnerability to social withdrawal, as shown in adult mice (Chen et al., 2020). Humans display prolonged myelination well beyond adolescence (Miller et al., 2012). Further experiments are needed to assess whether CT-179 produces clinically significant myelin toxicity in older children and adolescents. However, OLIG2 function in oligodendrocytes may be fundamentally different from OLIG2 function in tumour cells, where it modulates the chromatin landscape to activate a unique oncogenic program. CT-179 may therefore, specifically act on tumour cells without harming normal brain.

In summary, this study shows that inhibition of OLIG2-positive MB tumour cells in combination with RT significantly slows MB progression *in vivo.* CT-179, a novel, small molecule brain penetrate OLIG2 inhibitor, holds significant promise, particularly for the treatment of SHH-driven MB. CT-179 was recently awarded FDA Rare Paediatric Disease Designation for the treatment of MB in September 2020 paving the way for clinical testing of this effective OLIG2 inhibitor in children with SHH driven MB.

## Acknowledgements

Funding: This study has been supported by donations from the Children’s Hospital Foundation to the Children’s Brain Cancer Centre, The Sid Faithfull Group. TGJ is a Principal Research Fellow with the NHMRC. Additionally, this was supported by the NINDS (TD: F31NS120459; TRG: R01NS088219, R01NS102627, R01NS106227) and work by CL and MS was supported by the Carolina Partnership of the UNC Eshelman School of Pharmacy.

## Author Contributions

Y.L., T.D., C.L., T.G.J., T.R.G. and B.W.D. conceptualised the manuscript. Y.L., T.D., C.L., Z.C.B., C.O., U.B., R.C.J.D., T.R.G. and B.W.D. investigated the findings. Y.L., T.D., C.L., Z.C.B., C.O., and U.B. performed the analysis. M.M. and T.H., T.G.J., T.R.G. and G.S. provided the resource. Y.L., T.D., C.L., C.O., T.R.G., and B.W.D. performed the writing. B.W., M.P., T.R.G. and B.W.D. supervised the findings.

## Declaration of interests

G.S. is the CEO of Curtana Pharmaceuticals, Inc.

## STAR METHODS

### KEY RESOURCES TABLE

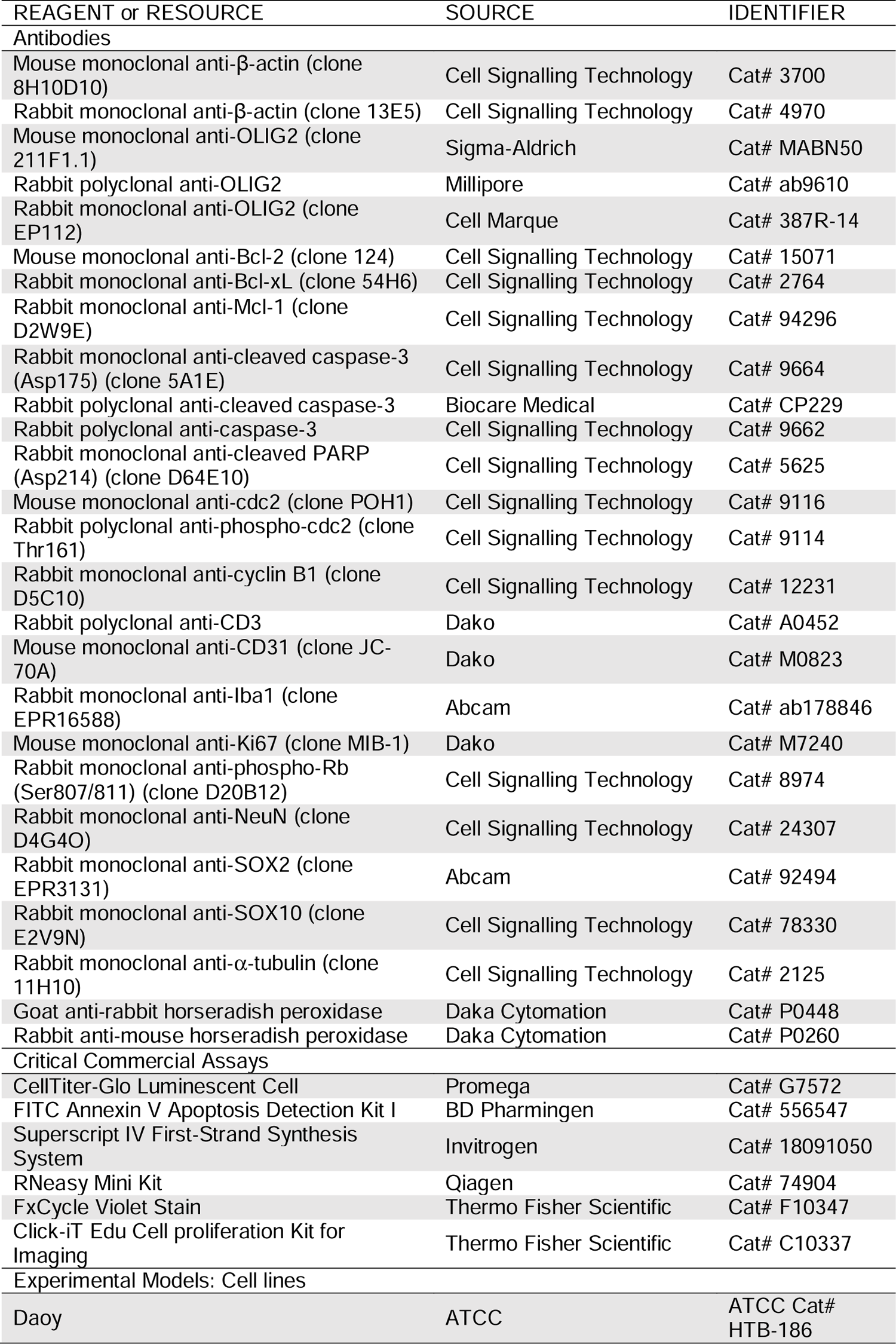
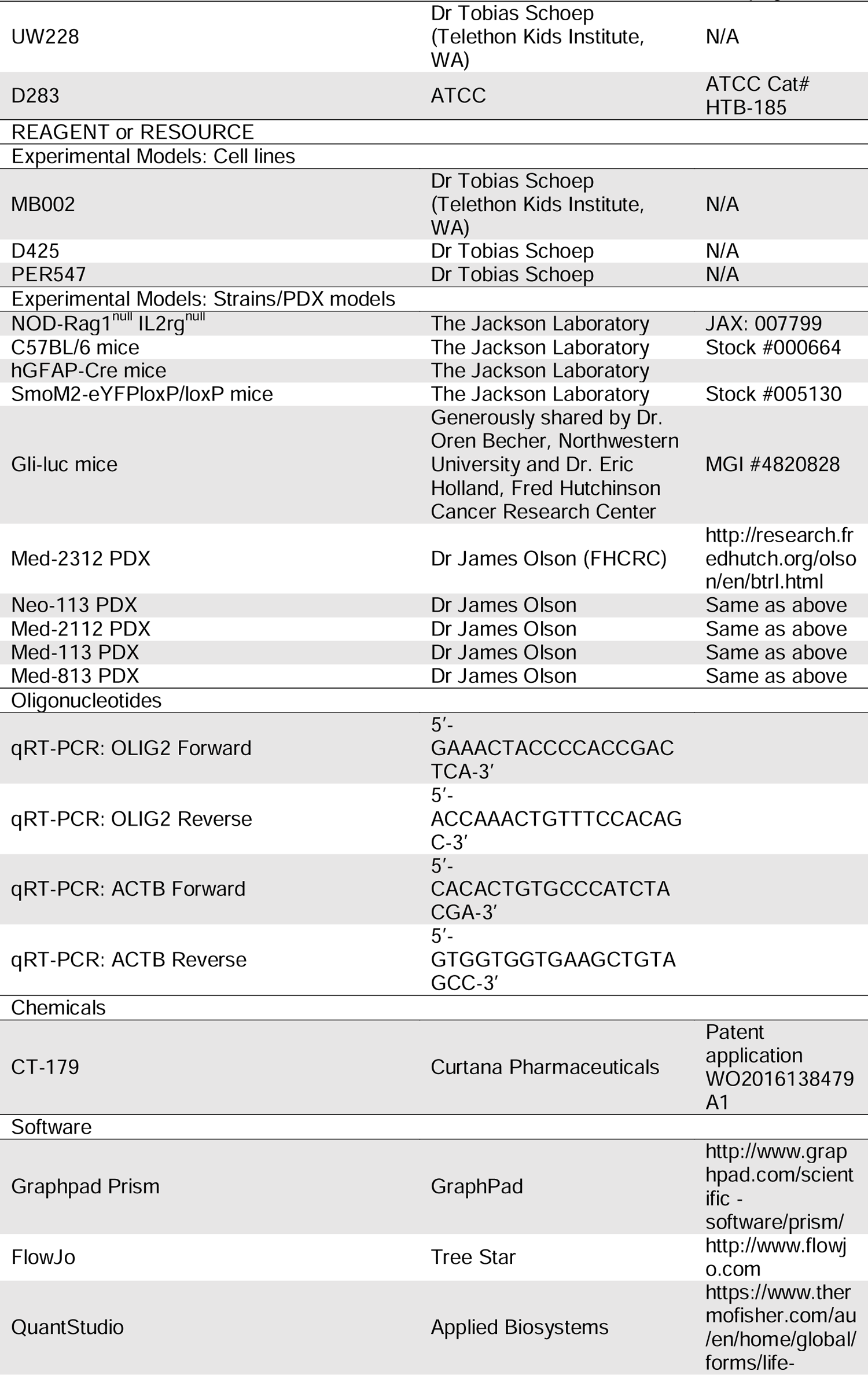
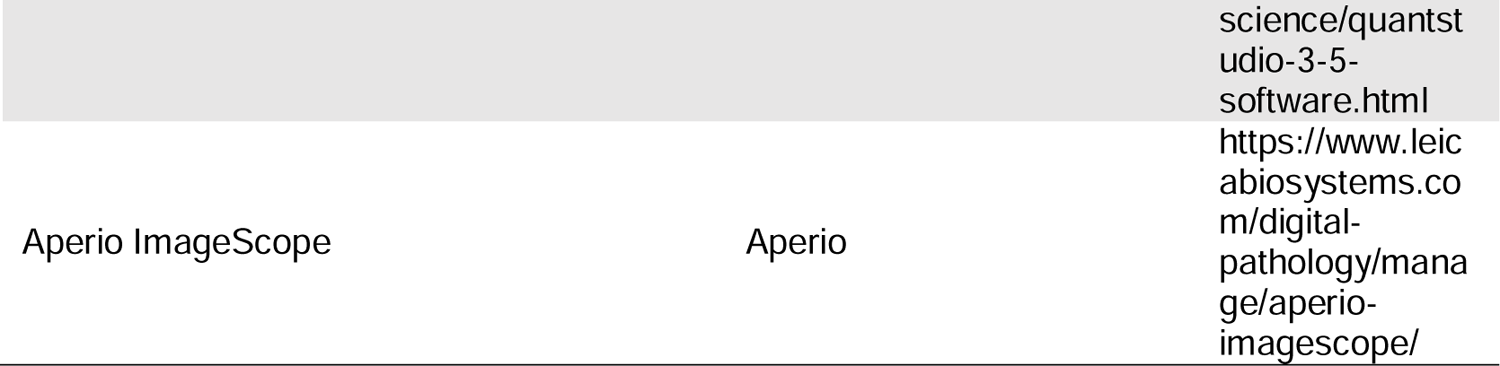

#### LEAD CONTACT

Further information and requests for resources and reagents should be directed to and will be fulfilled by the Lead Contact, Prof Bryan Day.

#### MATERIALS AVAILABILITY

All unique/stable reagents generated in this study are available from the Lead Contact with a completed Materials Transfer Agreement.

### Experimental Model and Subject details

#### Animal models

Human and animal experiment ethics approvals (P3420-A2102-601M and P2324-A1706-612M) were granted by the human ethics committee of the Queensland Institute of Medical Research (QIMR, Brisbane, QLD), Queensland Children’s Hospital (QCH) and Queensland Children’s Tumour Bank (QCTB). Female 6-7 week-old NOD-*Rag1^null^ IL2rg^null^* (abbreviated as NOD *rag* gamma, NRG) mice were used for toxicity and efficacy experiments. To assess the toxicity of CT-179, CT-179 (80 mg/kg) or saline was given to mice via oral gavage followed by RT (total 8 Gy, 2 Gy per fraction). Mice were monitored closely during the treatment. All mice were euthanised 4 days post treatment. Spleens and livers were collected and fixed in 10% neutral buffered formalin solution. Mouse blood was collected by cardiac puncture. The blood was analysed on a haematocytometer (Beckman Coulter).

To assess the efficacy of CT-179, mice received stereotactic-guided injection of live Daoy-luci cells (1 × 10^5^), Med-813-luci cells (3 × 10^5^) and Neo-113 filter tissue (FT). The stereotactic coordinates used to target the cerebellum, the orthotopic site of MB tumours, is as follows: 0.8 mm posterior to lambda, 1 mm lateral to the sagittal suture and at a depth of 2.5 mm. The Daoy and Med-813 models have been engineered to constitutively express the firefly luciferase gene (luci). Treatment started after tumour formation confirmed by bioluminescent imaging using the Xenogen IVIS^®^ Spectrum system (PerkinElmer, USA). CT-179 (50 mg/kg via IP, 75 mg/kg via oral gavage) or saline was given to mice followed by RT (total 8 Gy, 2 Gy per fraction). Tumour progression was monitored using the Xenogen IVIS^®^ Spectrum system. As per QIMR ethical guidelines, mice were sacrificed upon signs of tumour burden.

*G-Smo*-mice were maintained at UNC were handled in accordance with UNC IACUC protocols 19-098 and 21-011 and at QIMR under protocol 1572 and 2324. Specimens were examined by a neuropathologist to verify tumour type and grade.

#### Cell culture

MB cell lines Daoy, UW228, and D283 were obtained from the American Type Culture Collection (ATCC) and cultured in RPMI supplemented with 1% Penicillin-Streptomycin, 1% GlutaMax-I CTS (Gibco) and 10% fetal bovine serum (FBS) (Gibco). MB002, D425 and PER547 primary cell lines were kindly provided by Dr. Tobias Schoep (Telethon Kids Institute, WA). Med-2312 PDX, Neo-113 PDX, Med-2112 PDX, Med-113 PDX and Med-813 PDX were filter tissues purchased from the Olson Laboratory (https://research.fhcrc.org/olson/en/btrl.html). Cell lines derived from the corresponding PDX tissues were labelled as primary cell lines. MB-R201 PDX is a MB PDX model generated and established in our laboratory. MB-R203 is a primary MB cell line. Primary cell lines were derived from MB patient specimens and cultured either in serum-free media grown as spheres or on matrigel (Pollard et al., 2006). The serum-free media was prepared as previously described (D’Souza et al., 2020). Fresh surgically resected MB tissues were obtained from Queensland Children’s Tumour Bank. MB explant organoids (MBOs), R403 and R901, were generated by microdissection under a stereomicroscope (Zeiss) within a laminar flow biosafety cabinet. The MBOs were cultured in glioblastoma organoid (GBO) medium modified based on Jacob et al. (Jacob et al., 2020a). All cell lines were incubated at 37 °C under 5% CO_2_/95% humidified air atmosphere. All cell lines used in this study were authenticated by short-tandem repeat profile and tested for absence of mycoplasma contamination.

## METHOD DETAILS

### Reverse transcription and quantitative Real-Time PCR

Total RNA was extracted from cell lines and tissue samples using RNeasy Mini Kit (Qiagen, USA) according to the manufacturer’s instructions. First-strand cDNA was synthesized using Superscript IV (Invitrogen). Relative quantitation of gene expression was determined by real-time PCR which was carried out using SYBR^®^ Green PCR Master Mix (Applied Biosystems) following the manufacturer’s instructions. All reactions were performed in duplicate on an ABI Viia 7 (Applied Biosystems). Cycle thresholds (Ct) were determined and exported using QuantStudio™ software (Applied Biosystems). The mRNA transcripts levels of genes of interests were determined by the relative expression to the β tin reference gene. Primers used are listed in the Key resources table.

### Irradiation of cells

Cells were seeded into 96-well plates at 10,000 cells/well before irradiation in ^137^Cs source gamma rays to achieve 2 Gy (MDS Nordion Gammacell Irradiator).

### Cell viability assays

Cell proliferation was determined by a CellTiter 96^®^ Non-Radioactive Cell Proliferation Assay (Promega) kit. Absorbance at 490 nm was measured on a Biotek PowerWave (Biotek, USA) plate scanner to obtain raw data. Average absorbance from triplicate wells were plotted against concentration using GraphPad Prism software v.7.0 (GraphPad software). At the indicated time points, cells were stained using FITC-conjugated Annexin V and Annexin V binding buffer (BD Pharmingen^TM^) according to the manufacturer’s instructions. Annexin V or propidium iodide (PI)-positive cells were analysed using a Fortessa 5 Flow Cytometer (Becton Dickinson) and data was analysed using FlowJo^®^ software (Tree Star).

### CT-179 drug treatment

The synthesis of CT-179 is described in published patent application WO2016138479A1. To prepare the drug for oral gavage, CT-179 was dissolved in 0.5% (w/v) methylcellulose 4000/0.5% (v/v) Tween 80 in injection water. To prepare the drug for *in vitro* studies and for intraperitoneal (IP) injection, CT-179 was dissolved in injection water and stored at −80 °C.

### Cell cycle analysis

Cell cycle analysis was performed as descried previously (Linsley et al., 2007). Cells were fixed with 70% ice-cold ethanol and subsequently stained with PI and analysed using a Fortessa 5 Flow Cytometer (Becton Dickinson) and data was analysed using FlowJo^®^ software (Tree Star).

### Immunofluorescent staining

Cells were fixed with 4% paraformaldehyde (PFA) then quenched in blocking buffer (0.5% BSA), permeabilisation buffer (1 × PBS/0.3% Triton X-100) and then incubated with Hoechst and α-tubulin (Cell signalling), followed by incubation with Alexa Fluor^®^ 488 secondary antibody. Images were captured using a Zeiss 780-NLO confocal microscope (Zeiss) with a 40 × /1.4 Plan-Apochromat oil lens, and processed using Image J software.

Organoids were fixed in 4% paraformaldehyde (PFA)/ PBS, embedded in 1% low melting point agarose gel and, subsequently, embedded in paraffin. Organoid sections (4 µm) were probed with anti-SOX2 (Abcam), anti-Ki67 (Dako), CD3 (Dako), IBA1 (Abcam), and CD31 (Dako) either in single or in combination and subsequently stained with Mach 2 mouse/rabbit HRP or PE rabbit HRP. Sections were captured using a Zeiss 780-NLO confocal microscope (Zeiss) with a 40 × /1.4 Plan-Apochromat oil lens, and processed using Image J software.

For immunofluorescent staining of mouse MB after CT-179 treatment, brains with tumours were fixed, sectioned, stained and imaged using an Aperio Scanscope as previously described (Hwang et al., 2021).

### Immunoblot analysis

Whole cell protein lysate (60 μg/sample) was used. Samples were run on denaturing sodium dodecylsulfate-polyacrylamide gel electrophoresis gels (12%) before transferring onto Immuno-Blot™ polyvinylidene fluoride membranes (Bio-Rad). The reference protein β-actin was used as a loading control.

### In vivo pharmacokinetic studies of CT-179 in mice

The *in vivo* pharmacokinetic studies (EXT-240 and EXT-241) were performed under contract by Biodura Inc.

### Cell cycle analysis in *G-Smo* mice

Post-natal P10 *G-Smo* mice were injected IP with CT-179 (80 mg/kg) or saline control solution and tumours were harvested after 6 or 24 hours. EdU (40 mg/kg) was administered IP 30 minutes before harvest. Tumours were dissociated using the Worthington Papain Dissociation System Kit, then fixed for 15 minutes on ice and washed with FACS Wash buffer (2% FBS in PBS). Fixed cells were stained with fluorophore markers for DNA (FX Cycle stain, Thermo Fisher Scientific), EdU (Click-it Edu Kit, Thermo Fisher Scientific), and phospho-RB content (phospho-RB^Ser807/811^, Cell Signalling Technology) as previously described (Hwang et al., 2021). Stained cells were resuspended in sheath fluid and ran on LSR II flow cytometer for 50000 events at the UNC Flow Cytometry core, using appropriate compensation controls and analysed using FlowJo^®^ software (Tree Star).

### Immunohistochemistry

#### Tissues and organoids

Tissue samples were fixed in 10% neutral buffered formalin and embedded in paraffin and, subsequently, stained with haematoxylin and eosin (H&E). IHC on G-Smo mice was as previously described (Malawsky et al., 2021). Antigen retrieval was performed using a pH 9.0 Tris-EDTA buffer. Tissue sections (4 µm) were probed with anti-OLIG2 (Millipore) antibody overnight and subsequently stained with Mach Polymer HRP (Biocare Medical). Organoid sections (4 µm) were probed with anti-cleaved caspase 3 (Biocare Medical) in 10% goat serum for 2 hours at room temperature and subsequently stained with Mach 1 Universal Polymer HRP. Signals were developed in 3,3′-Diaminobenzidine (DAB). Sections were counterstained in Haematoxylin, dehydrated and mounted.

#### GEMM MBs

Brains including tumours from *G-Smo* mice were harvested, fixed in 4% paraformaldehyde for 48 hours, and embedded in paraffin at the UNC Centre for Gastrointestinal Biology and Disease Histology core. Sections were deparaffinised, and antigen retrieval was performed using a low-pH citric acid-based buffer. Staining was performed and stained slides were digitally scanned using the Leica Biosystems Aperio ImageScope software (12.3.3) by the UNC Translational Pathology Laboratory, as in prior studies (Malawsky et al., 2021). The primary antibodies used were anti-NeuN (Cell Signalling Technology) anti-OLIG2 (Cell Marque) and anti-SOX10 (Cell Signalling Technology).

## QUANTIFICATION AND STATISTICAL ANALYSIS

*In vitro* experiments - all data represent the means ± standard deviation (SD). Experiments were performed in three biological replicates. Where appropriate, two-tailed Student’s *t*-test or ANOVA were used to determine the probability of difference (*p < 0.05, ** p < 0.01, ***p < 0.001, ****p < 0.0001). *In vivo* experiments - A log-rank (Mantel-Cox) test was used to determine the survival significance of survival differences between treatment groups (*p < 0.05, ** p < 0.01, ***p < 0.001, ****p < 0.0001). Kaplan-Meier survival curves were generated using GraphPad Prism v. 7.0 software (GraphPad software).

