## Supplementary information for "Targeting OLIG2 increases therapeutic responses in SHH medulloblastoma mouse models and patient-derived medulloblastoma organoids"

**Table S1.** CT-179 or Radiation regimens used in *in vivo* testing

|  | **Control** | **CT-179 (C)** | | | | | **Radiation (R)** | | | | |
| --- | --- | --- | --- | --- | --- | --- | --- | --- | --- | --- | --- |
| **Day** | **n=16** | **#1 (n=6)** | | **#2 (n=7)** | **#3 (n=11)** | | **#1 (n=11)** | | **#2 (n=7)** | | **#3 (n=4)** |
| **10** | No treatment | 80 mg/kg | | 100 mg/kg | 80 mg/kg | | X | | X | | 2 Gy |
| **11** |  | X | | X | X | | X | | X | | 2 Gy |
| **12** |  | X | | X | 80 mg/kg | | X | | X | | 2 Gy |
| **13** |  | 80 mg/kg | | 100 mg/kg | X | | X | | X | | 2 Gy |
| **14** |  | X | | X | 80 mg/kg | | 0.5 Gy | | 1 Gy | | 2 Gy |
| **15** |  | X | | X | X | | 0.5 Gy | | 1 Gy | | X |
| **16** |  | 80 mg/kg | | 100 mg/kg | 80 mg/kg | | 0.5 Gy | | 1 Gy | | X |
| **17** |  | X | | X | X | | X | | 1 Gy | | X |
| **18** |  | X | | X | 80 mg/kg | | X | | X | | X |
| **19** |  | 80 mg/kg | | 100 mg/kg | X | | X | | X | | X |
| **20** |  | X | | X | 80 mg/kg | | X | | X | | X |
| **21** |  | X | | X | X | | X | | X | | X |
| **~35** |  |  |  | | | *  *  * |  |  | |  | |
| **Median survival** | **15.5** | **19.5** | | **20.5** | **21** | | **19** | | **16** | | **16** |

**Table S2.** Radiation + CT-179 regimens used in *in vivo* testing

|  | **Radiation + CT179 (C+R)** | | | | | | | |
| --- | --- | --- | --- | --- | --- | --- | --- | --- |
| **Day** | **#1 (n=4)** | | **#2 (n=4)** | **#3 (n=5)** | | **#4 (n=8)** | **#5 (n=4)** | **#6 (n=10)** |
| **10** | C+R  80 mg/kg+2 Gy | | C+R  80 mg/kg+1 Gy | C+R  80 mg/kg+1 Gy | | C 80 mg/kg | C 80 mg/kg | C 80 mg/kg |
| **11** | R 2Gy | | R 1Gy | X | | X | R 1Gy | X |
| **12** | C+R  80 mg/kg+2 Gy | | C+R  80 mg/kg+1 Gy | C+R  80 mg/kg+1 Gy | | C 80 mg/kg | C 80 mg/kg | C 80 mg/kg |
| **13** | R 2Gy | | R 1Gy | X | | X | R 1Gy | X |
| **14** | C+R  80 mg/kg+2 Gy | | C+R  80 mg/kg+1 Gy | C+R  80 mg/kg+1 Gy | | C+R  80 mg/kg+1 Gy | C 80 mg/kg | C+R  80 mg/kg+0.5 Gy |
| **15** | X | | X | X | | R 1Gy | R 1Gy | R 0.5Gy |
| **16** | C 80 mg/kg | | C 80 mg/kg | C+R  80 mg/kg+1 Gy | | C+R  80 mg/kg+1 Gy | C 80 mg/kg | C+R  80 mg/kg+0.5 Gy |
| **17** | X | | X | X | | R 1Gy | R 1Gy | X |
| **18** | C 80 mg/kg | | C 80 mg/kg | C+R  80 mg/kg+1 Gy | | C 80 mg/kg | C 80 mg/kg | C 80 mg/kg |
| **19** | X | | X | X | | X | R 1Gy | X |
| **20** | C 80 mg/kg | | C 80 mg/kg | C 80 mg/kg | | C 80 mg/kg | C 80 mg/kg | C 80 mg/kg |
| **21** | X | | X | X | | X | X | X |
| **~35** |  |  | | | *  *  * | |  |  |
| **Median survival** | **15** | | **19** | **18** | | **19** | **15** | **25.5** |

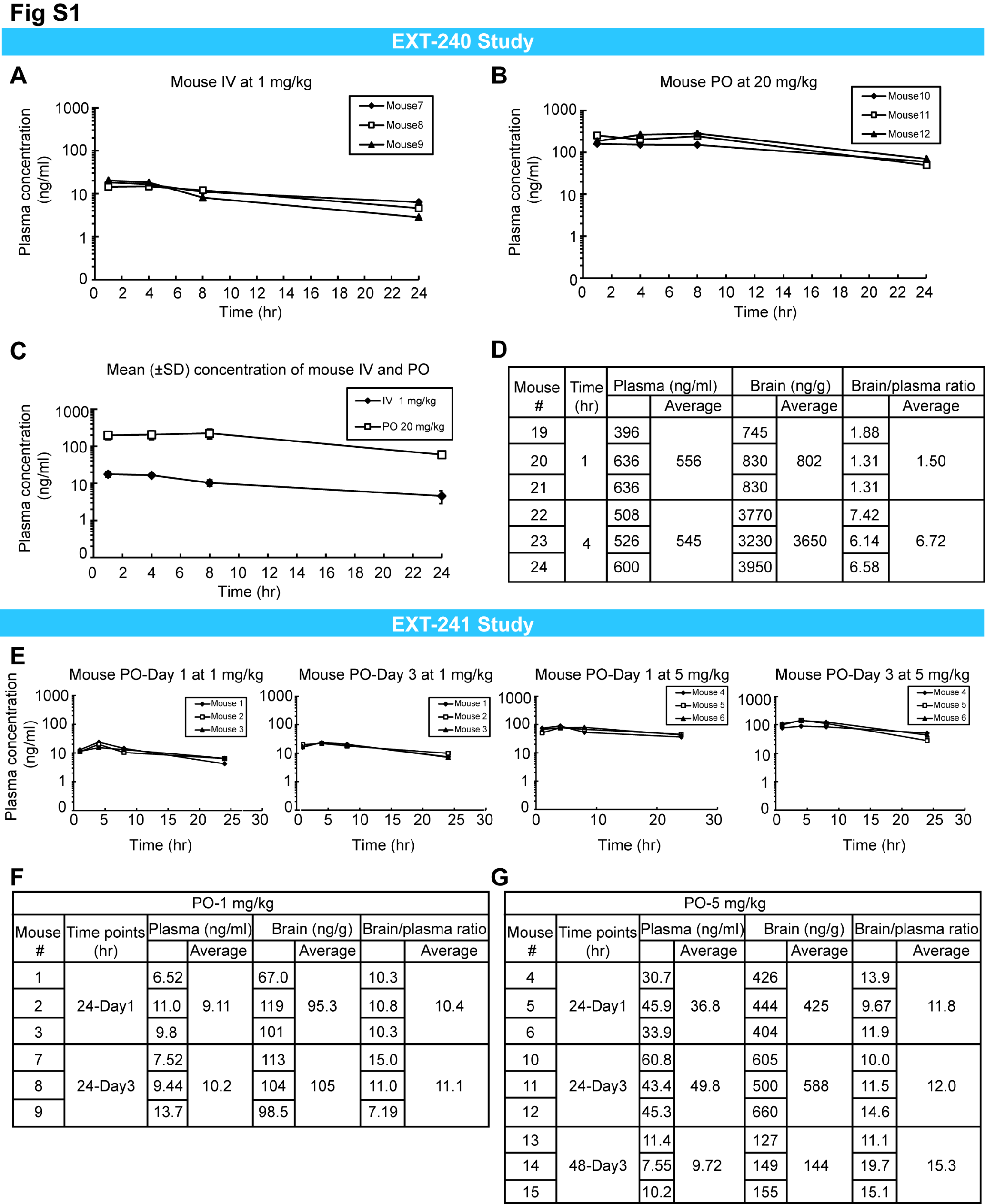

Supplementary Figure 1. CT-179 *in vivo* pharmacokinetic studies EXT-240 and EXT-241 in C57BL/J6 mice.

(A) Individual mouse data for intravenous (IV) CT-179 at 1 mg/kg.

(B) *In vivo* exposure results for CT-179 following an oral dose (per os, PO) of 20 mg/kg.

(C) A comparison of intravenous and oral dose groups.

(D) Summary of data for plasma and brain exposure at 20 mg/kg dose.

(E) Exposure results for CT-179 in mouse day 1 and day 3 following oral doses of 1 and 5 mg/kg.

(F) Summary data for plasma and brain exposure at 1 mg/kg oral dose.

(G) Summary data for plasma and brain exposure at 5 mg/kg oral dose.

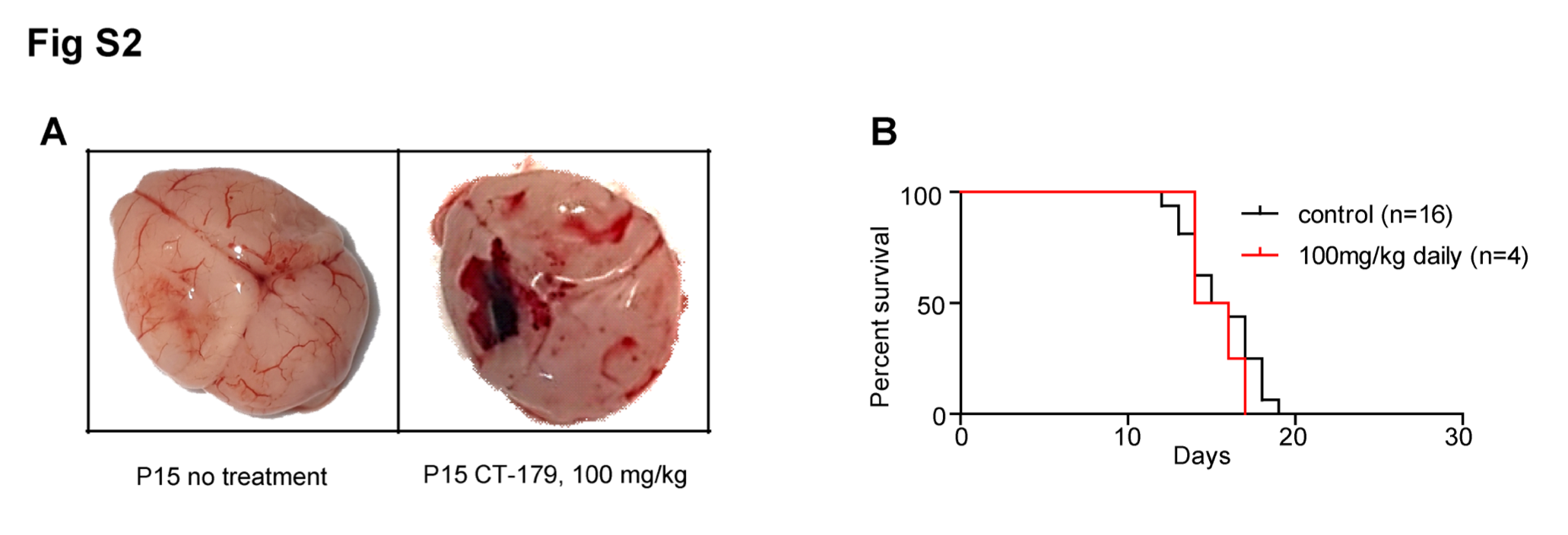

Supplementary Figure 2. Daily administration with CT-179 is met with toxicity limitations.

(A) Representative brains showing typical examples for haemorrhage after 100 mg/kg CT-179 treatment in *G-Smo* mice.

(B) Kaplan-Meier survival curve of *G-Smo* mice treated with control and CT-179 daily (100 mg/kg).

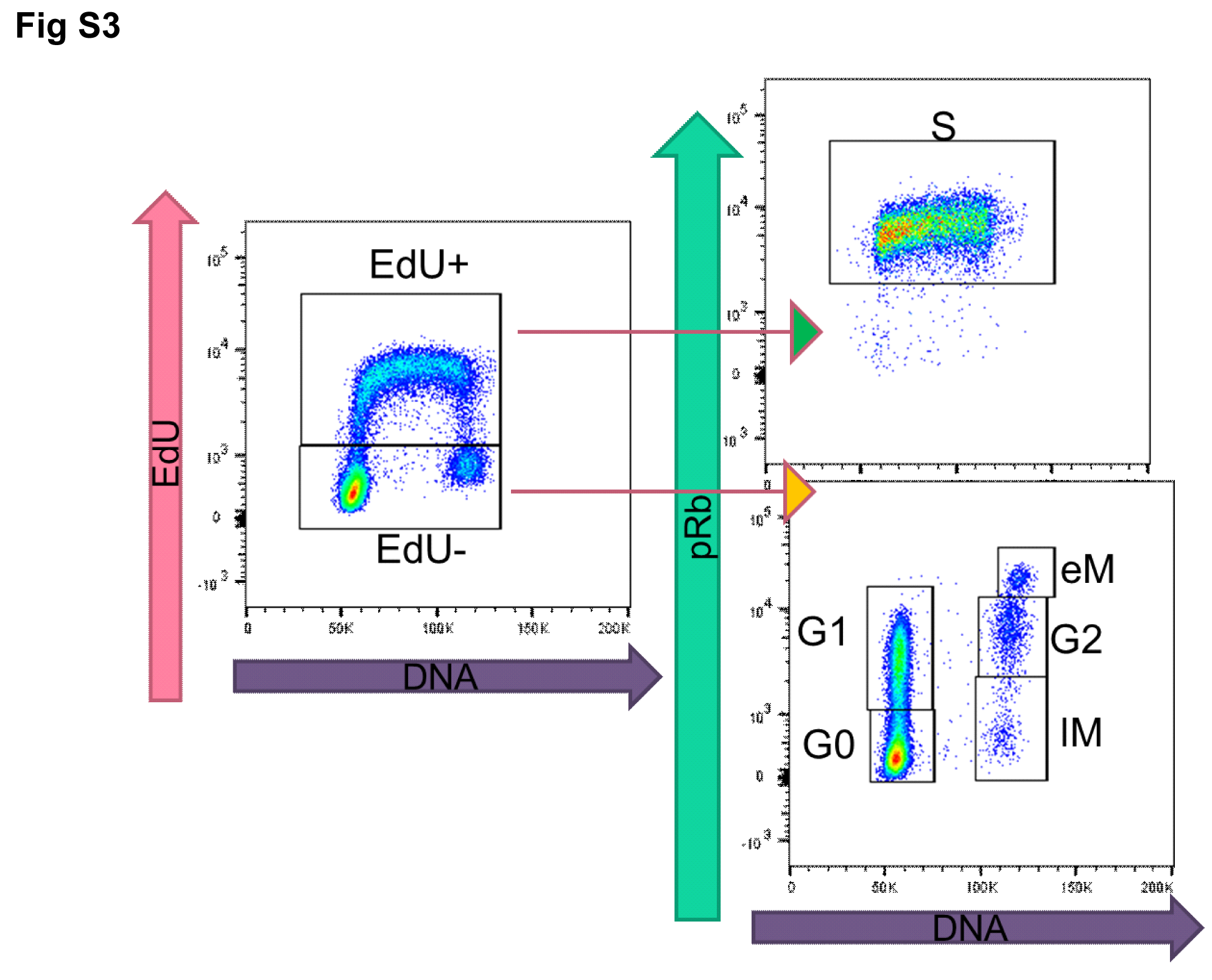

Supplementary Figure 3. Flow cytometry measurement of cell cycle dynamics in tumour from *G-Smo* mice.

Flow cytometry cell-cycle gating strategy was used to quantify dissociated tumour cells at G_0_, G_1_, S, and G_2_/M phases.

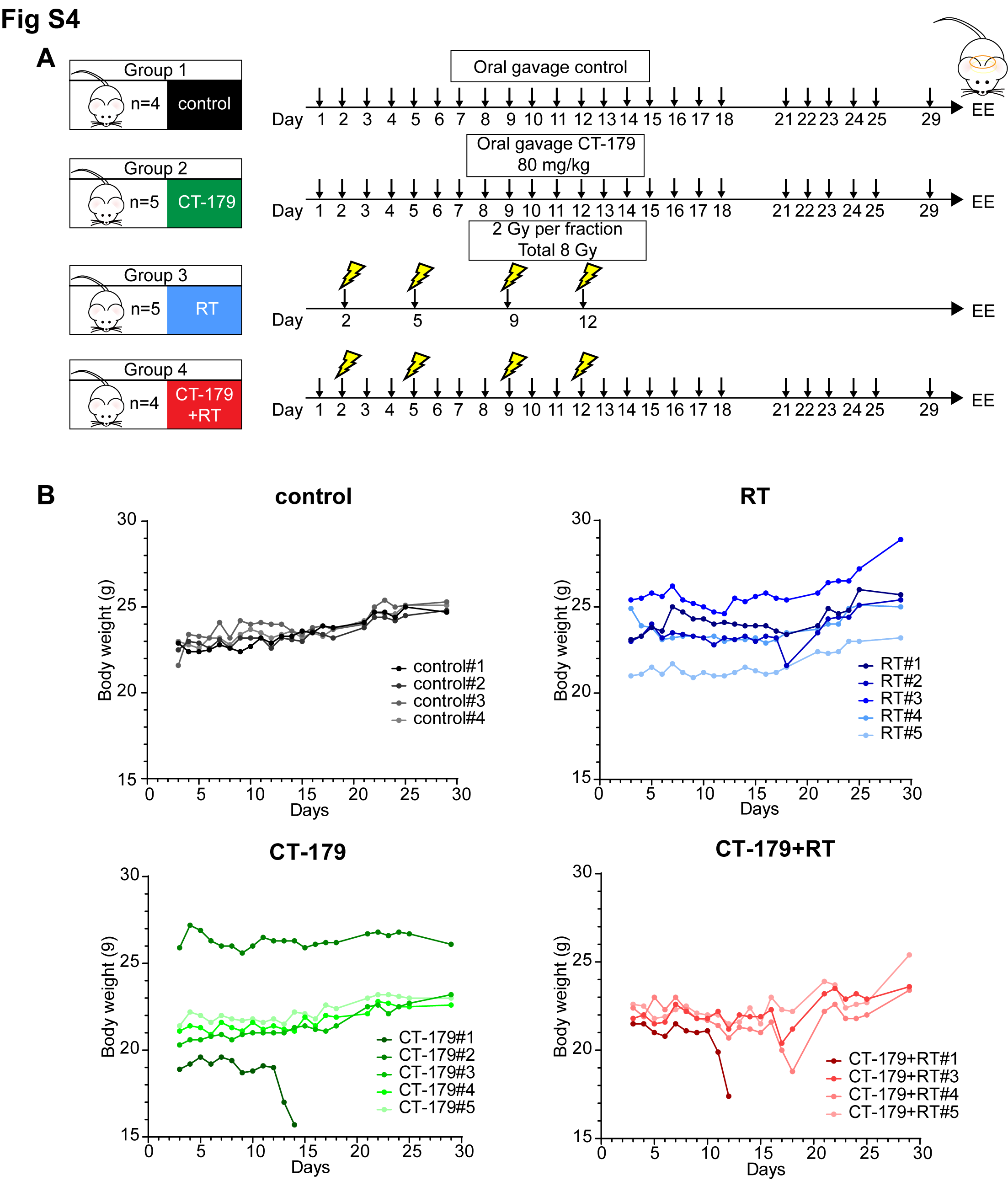

Supplementary Figure 4. Toxicity study in healthy NRG mice.

(A) Toxicity study treatment plans used in healthy NRG mice. Mice were assigned into 4 treatment groups, including vehicle (n=4), CT-179 (n=5), RT (n=5) and combination (n=4). Mice were euthanised on day 30 post commencement of treatment.

(B) Body weight of mice throughout the treatments and post treatments.

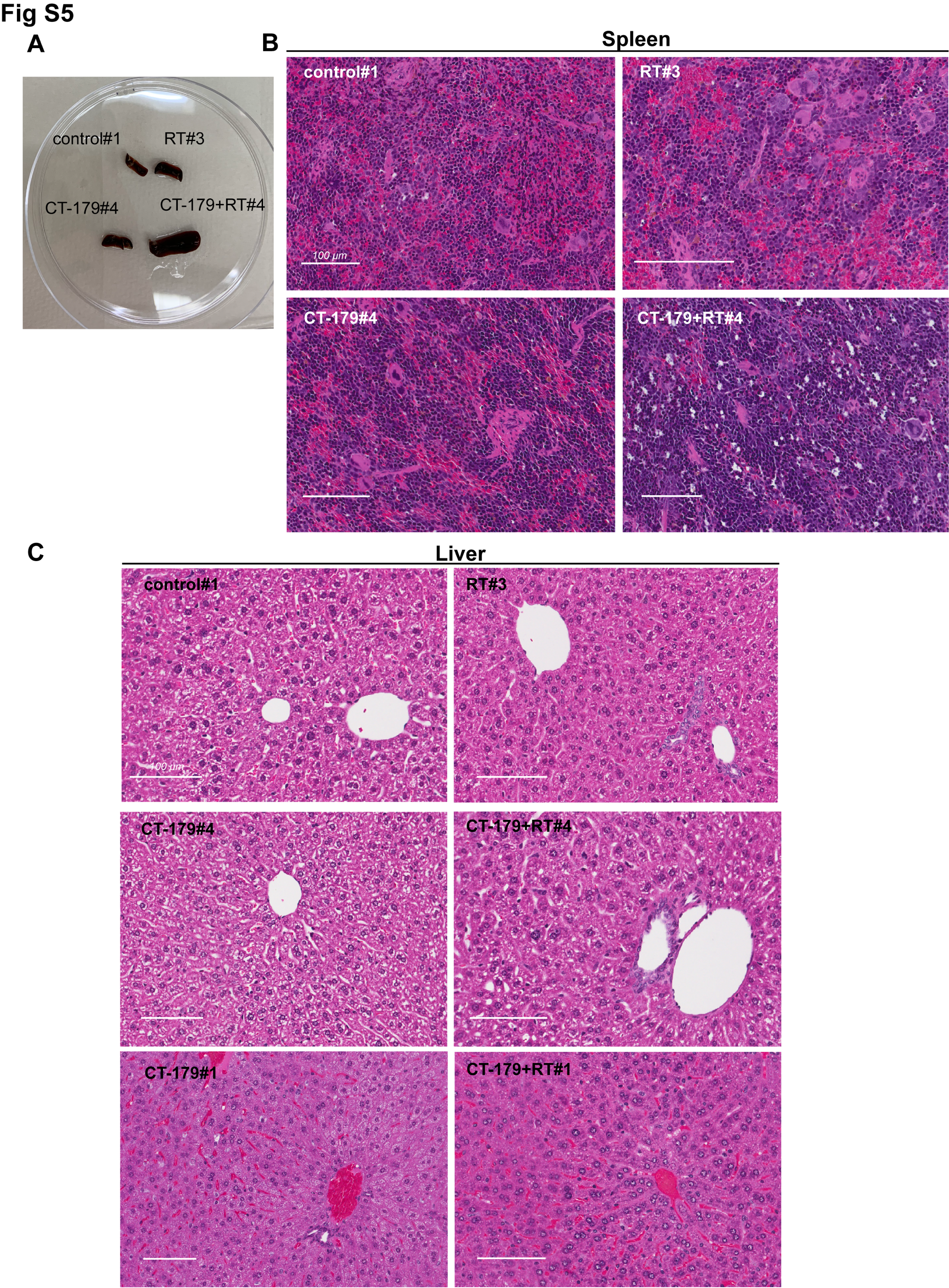

Supplementary Figure 5. Toxicities in spleen and liver found in healthy NRG mice post treatment.

(A) Spleens on day 30 post commencement of treatment. The spleen from mouse treated with CT-179 in combination with RT was significant enlarged.

(B) The histopathology of the spleen. Light microscope histological tissue slides of spleen from mice euthanised on day 30 post commencement of treatment. Control spleen section showing normal splenic architecture. Mouse treated with RT (RT#3) has ill-defined spleen section with diffused white pulp. Distorted lymphoid architecture and giant macrophages, presence of giant macrophages can be seen from CT-179 treated spleen. Presence of granular leukocytes in between lymphocytes in lymphoid follicles besides giant macrophages can be observed in mouse treated with combination treatment.

(C) Histopathology of the mouse liver post different treatment. Control, RT, CT179#4 and CT-179+RT#4 liver sections showing normal hepatic architecture, whereas livers from CT-179#1 and CT-179+RT#1 presented with haemorrhage features. The mice CT-179#1 and CT-179+RT#1 died in the middle of the treatment.

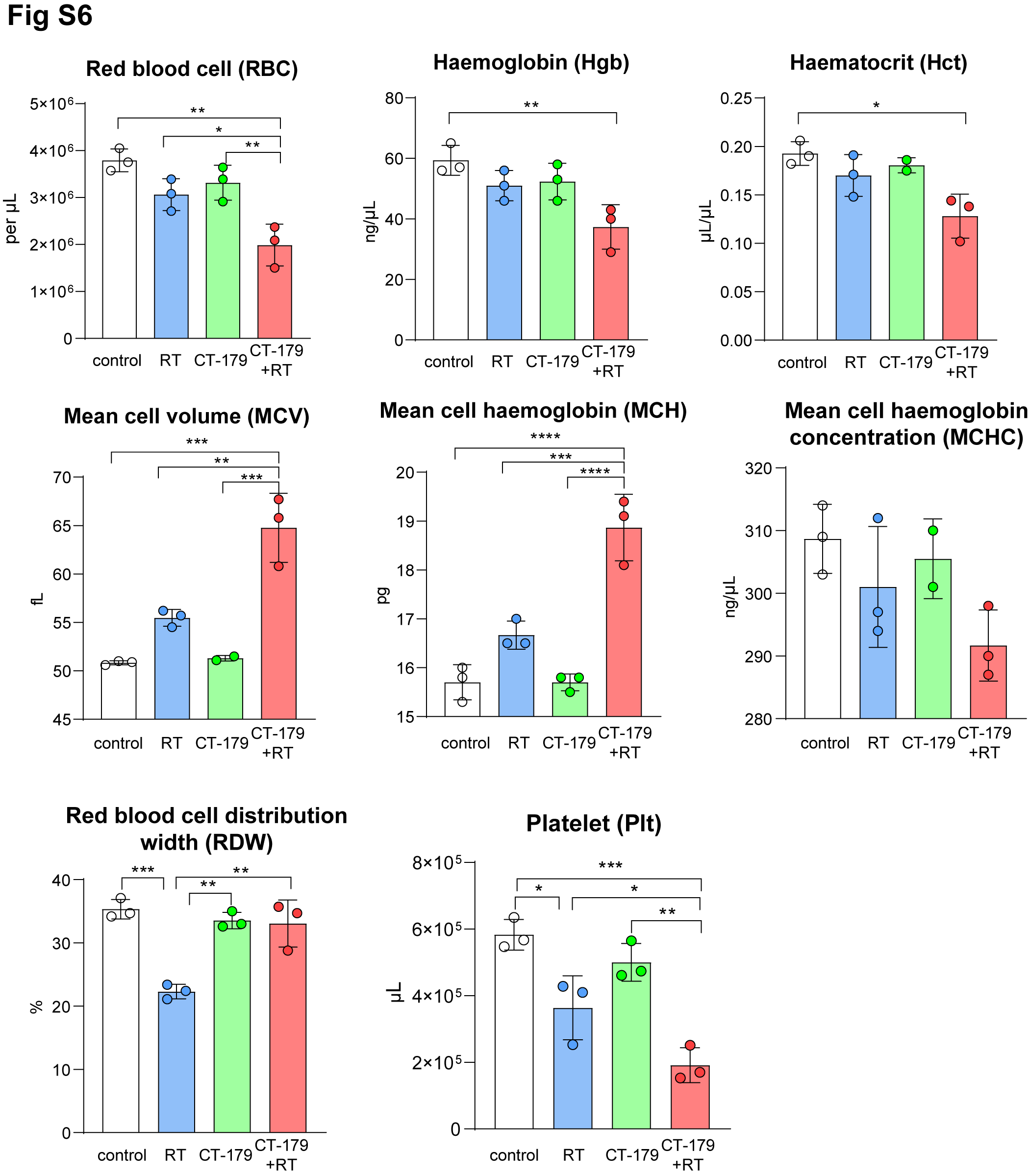

Supplementary Figure 6. Reduction of RBC and Plt in mice euthanised on day 30 post commencement of combination treatment.

Full blood count (exclude white blood cells) from mice euthanised on day 30 post commencement of treatment. Bar graphs are representing the red blood cell (RBC), haemoglobin (Hgb), haematocrit (Hct), mean cell volume (MCV), mean cell haemoglobin (MCH), mean cell haemoglobin concentration (MCHC), red blood cell distribution width (RDW) and platelet (Plt). RBC and Plt were significant lower when compared to the other treatment groups. Graphs display data from each individual animal from three independent experiments with each n=3. Statistical significance was determined using Student’s t test (means ± SD, n=3, *p < 0.05, **p < 0.01, **p < 0.001, ****p < 0.0001).

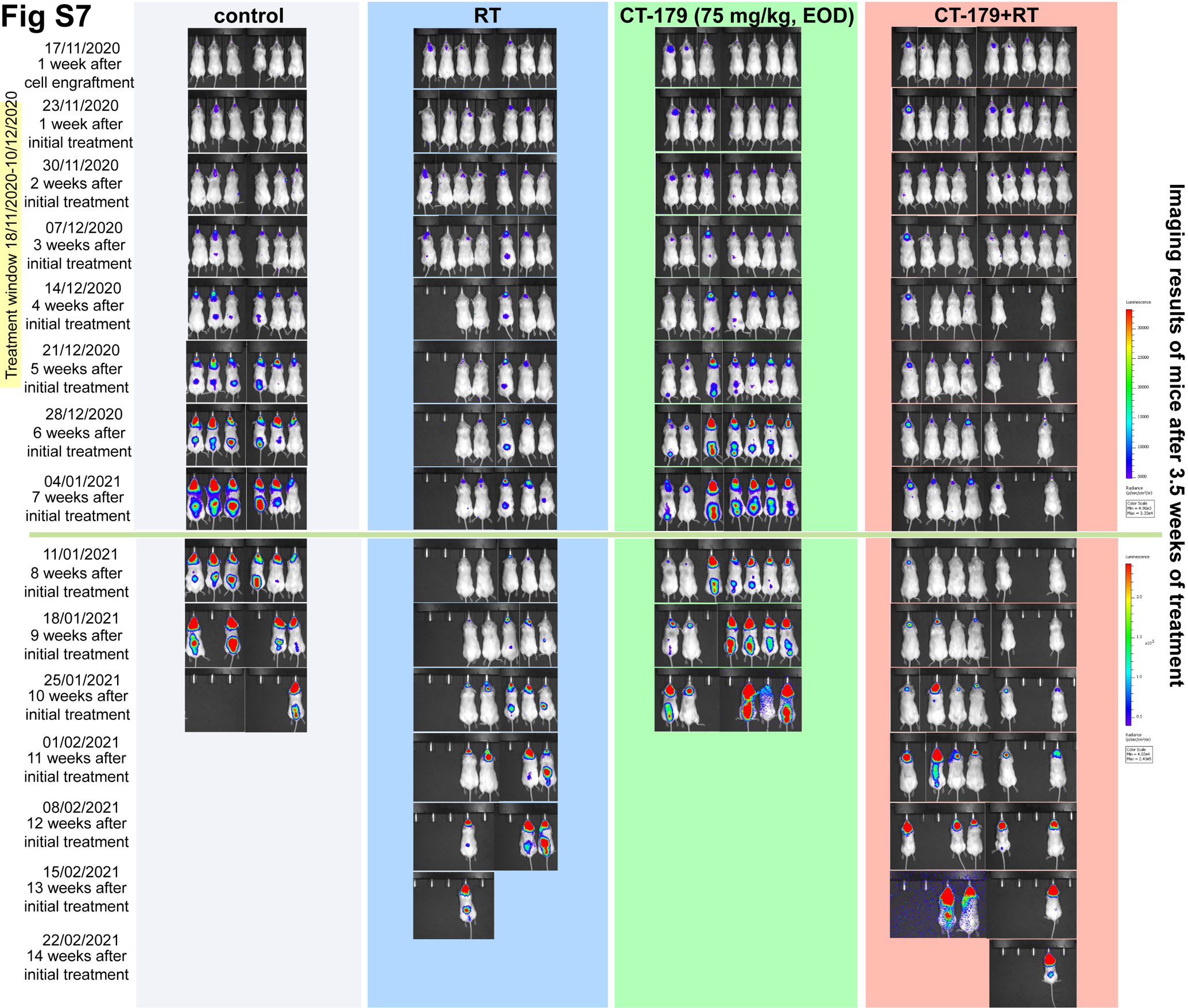

Supplementary Figure 7. Imaging results of Med813-luci model before and after 3.5 weeks of treatment.

Tumour formation was confirmed one week after cell engraftment using Xenogen. Tumour size was monitored throughout the treatments and after the treatments were completed. The graph displays mice with bioluminescent signals from 4 treatment groups, which are Vehicle control (n=6), Radiotherapy (RT, n=7), CT-179 (n=7) and combination treatment (n=9). Two mice from RT and three mice from combination group died from toxicity. Overall, mice in combination treatment group had prolonged survival followed by RT alone, CT-179 alone and vehicle. Tumour shrinkage was found in mice from CT-179 alone 4 weeks after the initial treatment. Similar results were observed in the combination treatment group. Most strikingly, we have seen less spinal metastasis in the mice from combination treatment group.
